# Disruption of interareal control during propofol anesthesia

**DOI:** 10.64898/2026.07.28.741350

**Authors:** Adam J. Eisen, André M. Bastos, Jacob A. Donoghue, Scott L. Brincat, Emery N. Brown, Ila R. Fiete, Earl K. Miller

**Author notes:** I.R.F. and E.K.M. are co-senior authors.

## Abstract

Anesthetic-induced unconsciousness may arise partly from a change in how brain areas can manipulate each other’s activity. To quantify this change, we use control-theoretic tools that can precisely characterize how easily subsystems in complex networks can control each other. These tools rely on the Jacobian of the dynamics, an object that fully specifies how inputs to a function affect the outputs. We built on JacobianODE, a method for data-driven Jacobian learning, to enable its application to neural recordings. We developed a deep learning framework that recovers directional, nonlinear control from partially observed multi-area recordings by combining delay-coordinate embedding, a volume-preserving invertible encoder, and latent dynamics via JacobianODE. We validated our framework on the Lorenz system and a partially observed working-memory recurrent neural network. We then applied it to local field potential recordings from posterior parietal (PPC), superior temporal gyrus (STG), frontal eye fields (FEF), and ventrolateral prefrontal cortex (vlPFC) in two non-human primates, comparing wakefulness with propofol anesthesia. Anesthesia pervasively reduced the ability of areas to control each other, both in terms of driving towards novel states and stabilizing along existing trajectories. This change in ease of control was driven by a decrease in the magnitude of interareal coupling. Directional ease of driving control from PPC to vlPFC and FEF to vlPFC was increased under anesthesia, providing a potential mechanism for paradoxical excitation observed during propofol infusion. Together, these results recast anesthetic unconsciousness as a directional breakdown of cortical control.

## 1 Introduction

Neural computation is supported by the dynamic, directional interactions between brain areas. Cross-area interaction underwrites a remarkably broad swath of neural function:^1,2^ it mediates the selective routing of attention,^3–11^ supports decision making,^12,13^ sustains working memory,^14,15^ binds features into coherent percepts,^16,17^ coordinates motor output,^18–22^ and underlies learning and memory.^23–27^ It is therefore unsurprising that interareal interactions are thought to underlie conscious experience itself.^28–35^

General anesthetics produce a profound, reversible loss of consciousness. They thus present a promising setting for the study of consciousness in the brain.^36–41^ By comparing anesthetic-induced changes in neural dynamics between conscious and unconscious states, and identifying robustly altered interareal interactions, we aim to gain insight into the neural mechanisms underlying consciousness. Propofol, a GABA_A_ (*γ*-aminobutyric acid type A) receptor agonist, changes the balance between excitation and inhibition.^42–48^ It has been shown to produce robust disruptions to cortical connectivity and integration.^37,45,49–64^

Control theory formalizes how external inputs can be coordinated with a system’s intrinsic dynamics in order to elicit desired behaviors, with applications spanning engineering, robotics, and biology (Figure 1A). Given the dynamic nature of cortical computation, control theory is a natural lens through which to interpret interactions in the brain (Figure 1B).^19,65–71^

**Figure 1.**
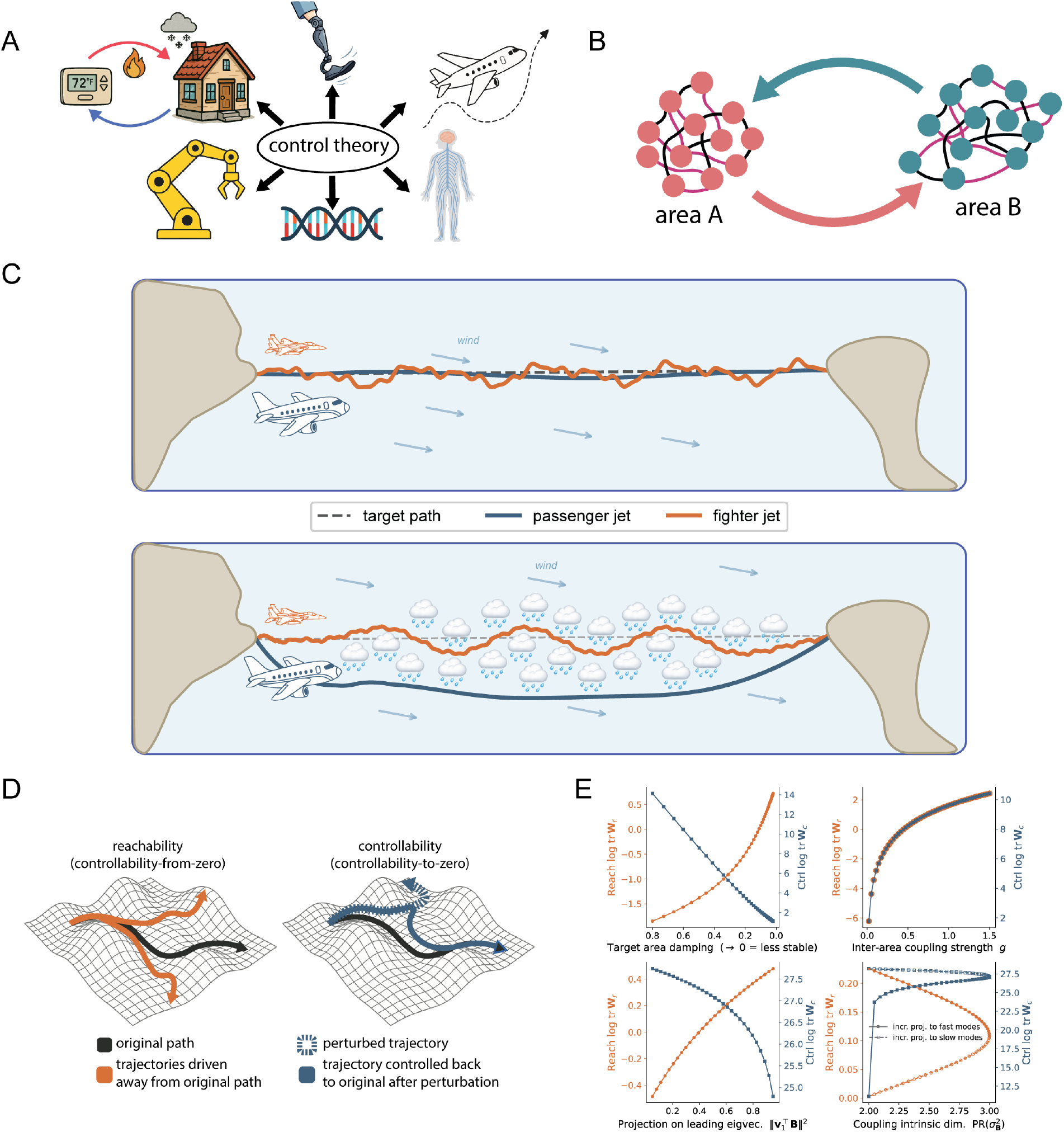
A control theoretic framing of interareal control. (A) Control problems share a common formal structure across engineering, biology, and brain networks. (B) Bidirectional control interactions between two areas. (C) Conceptual illustration of reachability and controllability interacting with dynamic stability. (D) Abstract illustration of reachability and controllability. (E) The dependence of reachability and controllability on dynamic stability, coupling magnitude, coupling alignment, and coupling dimensionality in a synthetic two-area linearization.

Two complementary quantities formalize measurements of ease of control (Figure 1C–E). Along a reference trajectory, *reachability* measures how easily the target system can be driven away from its trajectory, while *controllability* measures how easily the target system can be driven back toward its trajectory.^72^ Both trade off against the target’s dynamic stability (Figure 1C). Consider a passenger jet and a fighter jet attempting to fly in a straight line. The passenger jet is more dynamically stable, meaning it more easily returns back to its original heading after being perturbed by disturbances such as turbulence. When the goal is to stay on course (controllability, Figure 1C, top and Figure 1D, right), the passenger jet can more easily be guided back to the straight line path in the face of disturbances. However, when the planes must deftly navigate around storm clouds (reachability, Figure 1C, bottom and Figure 1D, left), the fighter jet can more easily be driven to deviate from its original path and sharply steer around the obstacles.

Reachability and controllability thus have opposite relationships with the dynamic stability of the target system (Figure 1E, top left). As dynamic stability increases, controllability increases while reachability decreases. Coupling strength, on the other hand, is positively correlated with both reachability and controllability (Figure 1E, top right). Greater alignment of the coupling matrix with the leading eigenvector of the target area increases reachability (perturbations propagate along more unstable modes) but decreases controllability (harder to stabilize more unstable modes) (Figure 1E, bottom left). Accordingly, coupling intrinsic dimensionality can have varying effects, depending on whether the increased dimensionality distributes onto slower or faster modes of the target area (Figure 1E, bottom right). Thus reachability and controllability together depend intricately on a multitude of factors, including dynamic stability, coupling strength, as well as coupling dimensionality and the alignment of the inputs with the target system’s dynamics.

The lack of attentional control and awareness during anesthetic-induced unconsciousness suggests a dramatic change in control dynamics. Indeed, propofol and other general anesthetics have been shown using linear network control theory to increase the transition energy required to switch states from functional imaging data.^73^ While existing neural control analyses have leaned on linear formulations.^74–86^, the brain’s dynamics are fundamentally nonlinear, and the context-dependent forms of control they afford cannot be captured in their entirety by linear models, motivating a nonlinear treatment of interareal control.^87–91^

In this work, we leverage nonlinear control theory to characterize how interareal interactions reorganize under propofol anesthesia from intracortical depth electrodes recording local field potentials (LFPs). Our analysis uses the Jacobian of the dynamics, a matrix-valued function that captures the local interdependence of the dynamics of every component of the system. We reproduce findings from previous work on propofol anesthesia that found propofol induces destabilized dynamics.^92,93^ We then characterize how propofol dismantles both reachability and controllability in the brain, on top of demonstrating a reduction in the magnitude and dimensionality of local interareal coupling. Overall, our results suggest that propofol anesthesia disrupts consciousness through the disruption of interareal directional control dynamics.

## 2 Jacobian-based directional control estimation

We consider nonlinear dynamical systems in ℝ ^d^. The full system is defined by the dynamics 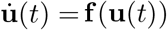. We assume that within this full system, there are *K* coupled subsystems, each of which is defined by the dynamics 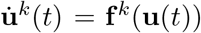, *k* = 1, …, *K*. The subsystem dimensionalities are *d*_k_, and 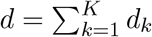. The system is then observed through a smooth map 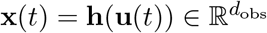 with 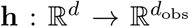 that projects the full state vector **u** onto a lower-dimensional subspace. We will assume that our observation of each subsystem depends only on the state of the subsystem itself, and not on the state of the other subsystems. That is, we can decompose the observation map as **x**(*t*) = **h**(**u**(*t*)) = [**h**^1^(**u**^1^(*t*)), **h**^2^(**u**^2^(*t*)), …, **h**^K^(**u**^K^(*t*))].

### 2.1 Jacobian-based directional control estimation in fully observed dynamical systems

We begin with the simplest case discussed in the original JacobianODE paper^87^, when **h** is the identity map, so **x**(*t*) = **u**(*t*). The Jacobian of the dynamics is a matrix-valued function **J**_**f**_ : R^d^ → R^d*×*d^ (henceforth, **J**) given by

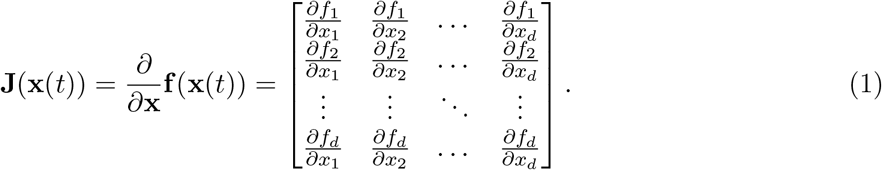

At each time *t*, the Jacobian captures how perturbations to the system will propagate. This recasts nonlinear dynamics as linear time-varying dynamics in the tangent space locally along trajectories (formally,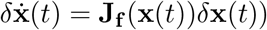 (Figure 2A, left).

**Figure 2.**
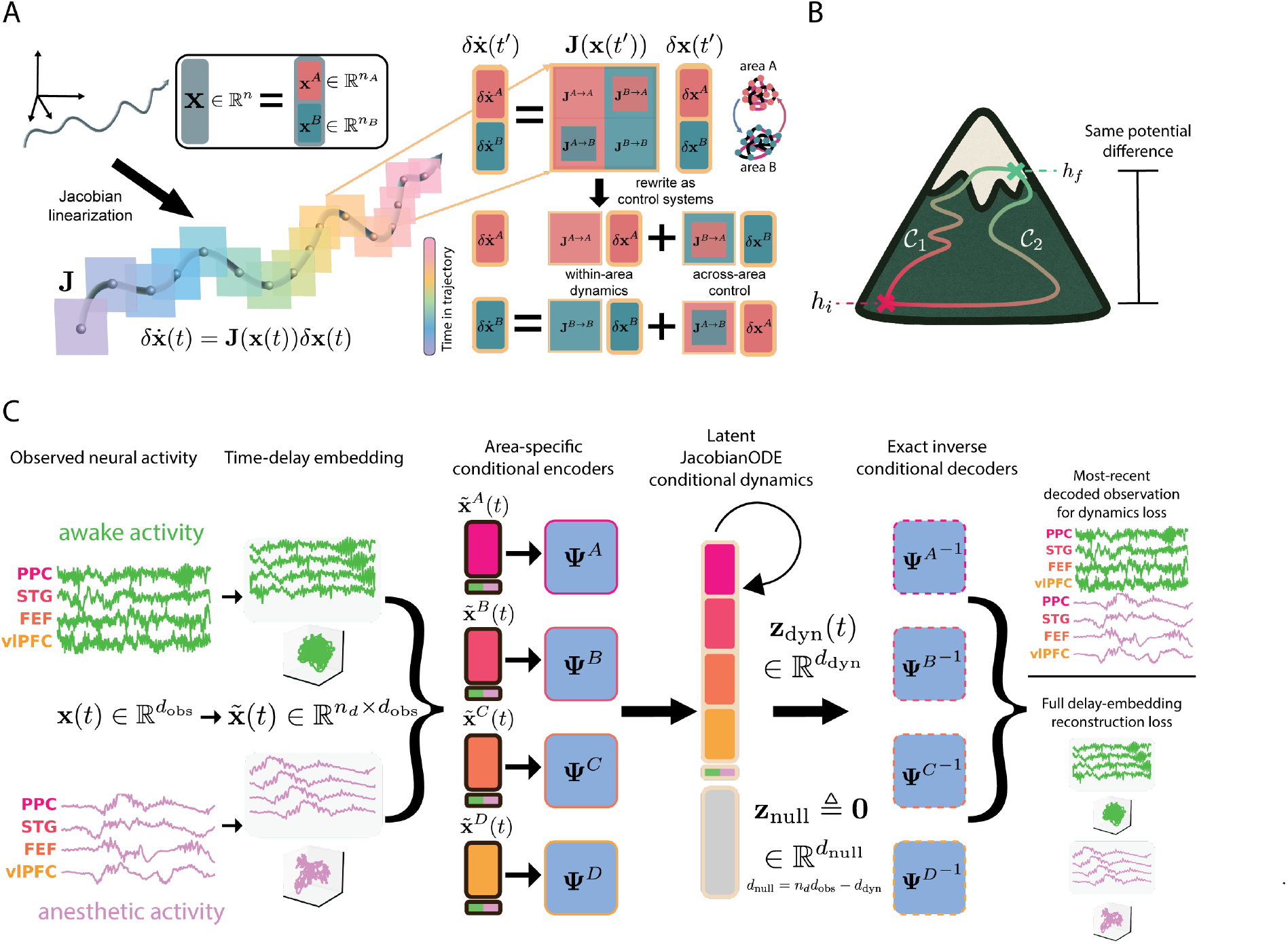
Latent JacobianODE pipeline for partially observed multi-area neural recordings. (A) Jacobians linearize dynamics along a reference trajectory, enabling the construction of pairwise interacting control systems. (B) Jacobians obey path independence, analogous to achieving the same potential energy difference taking two different paths up a mountain with the same endpoints. (C) The latent JacobianODE deep learning framework, demonstrated using the four-area neural recordings analyzed in this work.

To characterize control, consider the following formulation. As discussed in the original JacobianODE paper, the tangent space dynamics around the reference trajectory for subsystem *α* ∈ {1, …, *K*} are given by

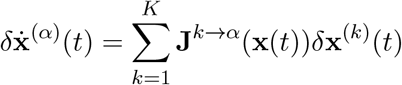

∈ { }

where 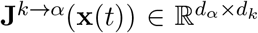 is the submatrix of **J**(**x**(*t*)) in which the columns correspond to sub-system *k* and the rows correspond to subsystem *α*. Now, for a given subsystem *β* ∈ {1, …, *K*} with *β* ≠ *α*, we wish to analyze the ease with which *β* can control *α* locally around the reference trajectory, without intervention from other subsystems. Discounting the interventions from other subsystems equates to setting *δ***x**^(k)^(*t*) = 0 for *k*≠ *α, β*, leaving the expression

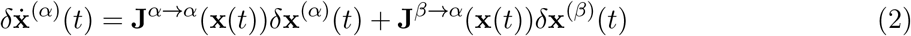

which quantifies the local dynamics of subsystem *α* with input from subsystem *β*, in the absence of perturbations from any other subsystem (Figure 2A, right).

To quantify the ease of control subsystem *β* can exert over subsystem *α*, we can use the reachability and controllability Gramians, defined in continuous time as

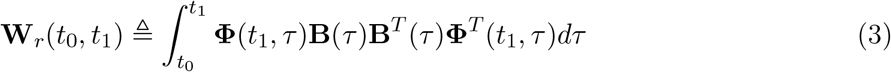

And

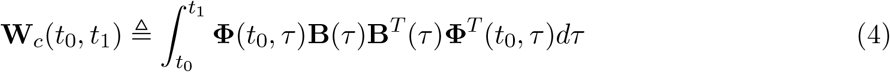

respectively. In these equations, **Φ** denotes the state-transition matrix of the intrinsic dynamics of subsystem *α* without any input (i.e., *δ***x**^(α)^(*t*) = **Φ**(*t, t*_0_)*δ***x**^(α)^(*t*_0_)) and *B* denotes the coupling Jacobian **J**^β*→*α^(**x**(*t*)).^94^

### 2.2 Deep Jacobian estimation: learning Jacobians from data

Given only observed trajectories 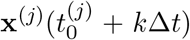, *k* = 0, 1, …, and no access to the underlying dynamics **f**, we estimate **J** directly by parameterizing it as a neural network **Ĵ**^θ^ with learnable parameters *θ*.^87^ The core observation is that each row of **J** is a conservative vector field (the gradient of a scalar potential), so its path integral between any two points is path-independent^95,96^: as in the work done climbing a mountain, the result depends only on the start and end points, not the route taken (Figure 2B). Formally,

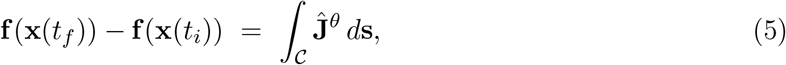

where is any piecewise-smooth curve joining **x**(*t*_i_) to **x**(*t*_f_). Together with a Jacobian-based parameterization of the reference value **f** (**x**(*t*_i_)) so that all gradients backpropagate through *θ* (see Appendix B.1), Equation 5 yields an estimate of the time derivative everywhere along the trajectory, which a standard ODE integrator advances to predict 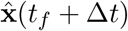.

#### Trajectory loss

Given an observed trajectory **x**(*t*_0_ + *k*Δ*t*), *k* = 0, …, *T* − 1, the trajectory reconstruction loss ℒ _traj_(*θ*; **x**) penalizes deviation between predicted and true states under an appropriate distance (typically mean squared error, i.e. MSE). Recursive predictions in chaotic or noisy systems are stabilized by generalized teacher forcing, which partially anchors the rollout to ground-truth states.^97^

#### Loop-closure loss

Trajectory loss only constrains **Ĵ**^θ^ along the direction of the flow, leaving the orthogonal tangent-space directions underdetermined. Exploiting conservativity once more, the path integral of **J** around any piecewise-smooth closed loop _loop_ must vanish (Figure 2C). The self-supervised loop-closure regularizer^98^,

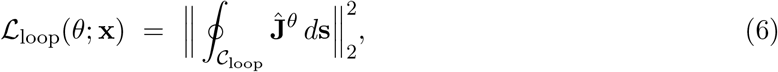

penalizes the deviation of this integral from zero. In practice, we form from concatenations of line segments between randomly-sampled data points, sampling diverse tangent-space directions on the data manifold while remaining cheap to compute.

#### Training objective

The total loss combines the two terms,

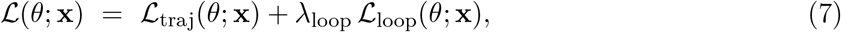

with *λ*_loop_ a tunable hyperparameter controlling the weight of the loop-closure regularizer.

### 2.3 A latent JacobianODE pipeline for partially observed dynamical systems

#### Delay embedding

To recover the full system dynamics from partial observations, we can leverage Takens’ delay embedding theorem. The theorem guarantees that by looking backwards in time at the history of the observed data, we can reconstruct a manifold that is diffeomorphic to the full system state space. Specifically, we form delay-embedded vectors 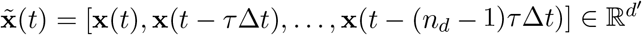, with *d*^*′*^ = *n*_d_*d*_obs_. Taking the intrinsic dimensionality of the system to be *d*^⋆^, Takens’ theorem guarantees that for generic **h** and *n*_d_ ≥ 2*d*^⋆^ + 1, the map 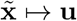 is a diffeomorphism from the embedded manifold onto the full system state space. We will again assume subsystem-specificity of the observation map, so that 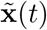 can be written as 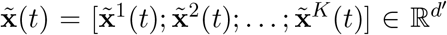, and the diffeomorphism is given by 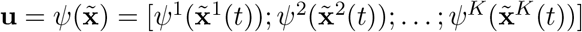.

**Figure S1.**
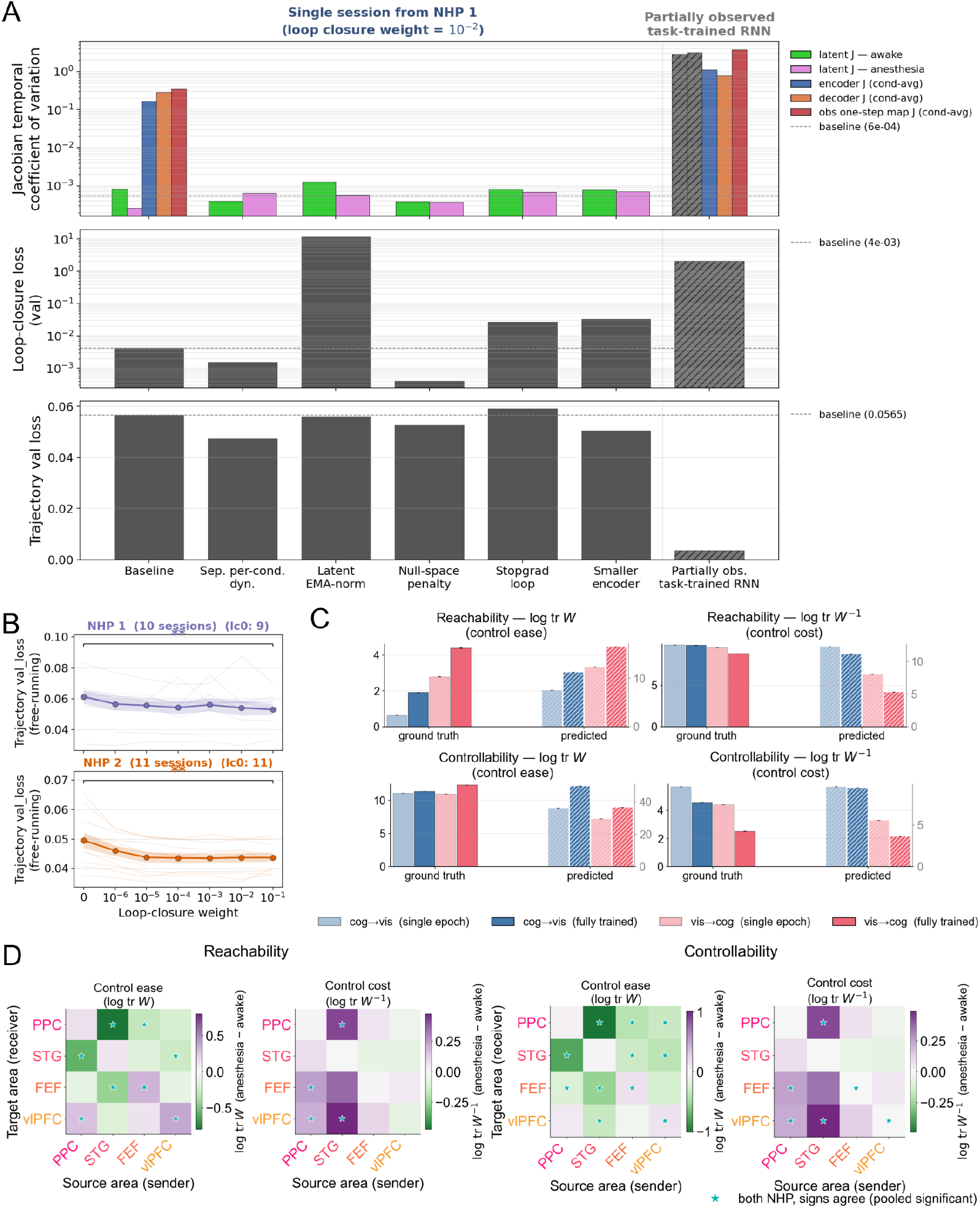
Linearity diagnostics and linear delay-embedded baseline. **(A)** Baseline with five anti-LTI interventions on a single session from NHP 1 (using loop-closure weight 10^−2^), with a partially observed task-trained RNN as nonlinear reference. (A, top) Per-condition latent Jacobian temporal coefficient of variation (CV), used as a measure of nonlinearity. The condition-averaged encoder (**J**_**Ψ**_), decoder (**J**_**Ψ***−*1_), and round-trip Jacobian CV area overlaid for the baseline and the RNN reference. (A, middle) Validation loop-closure loss. (A, bottom) Free-running trajectory validation loss. The NHP 1 variants cluster near the baseline on all three latent-Jacobian diagnostics. The encoder/decoder Jacobians, by contrast, are two to three orders of magnitude more state-dependent (Eq. 14), and the reference shows the genuinely nonlinear regime the architecture is capable of. **(B)** Best free-running trajectory validation loss vs. loop-closure weight *λ*_loop_, per NHP, across the cohort’s chosen-cell sweep grid (per-session lines + mean ± SEM). There is a monotone-non-decreasing trend in both NHPs, indicating improved fits with models that obey loop closure laws.. **(C)** Per-area linear delay-embedded baseline (Eqs. 15–16) applied to the partially observed task-trained RNN. Reachability and controllability gramian and inverse gramian log traces (log tr **W** and log tr **W**^*−*1^) for the visual → cognitive and cognitive → visual blocks at single-epoch and fully-trained checkpoints. Ground truth (left of each pair) vs. linear baseline (right). **(D)** The same linear baseline on the propofol cohort, with the four sign-agree pooled-Wilcoxon 4 × 4 Gramian Δ-heatmaps that mirror Figure 5A,B of the main text. The qualitative reorganization is preserved, with smaller effect sizes and more variability.

#### Diffeomorphic encoder

The latent version of the JacobianODE model builds on the insight from Takens’ theorem, and following previous work,^99^ leverages diffeomorphic encoders to learn a latent representation that is diffeomorphic to the delay embedding. Specifically, we delay embed the observed data, and construct an encoder that consists of coupling layers interleaved with orthogonal mixing layers. This results in an encoder that is both diffeomorphic and exactly invertible (Figure 2C, left). In practice, we use additive coupling layers for the coupling component, and Cayley-parameterized orthogonal mixers for the mixing component.^100–102^ Formally, an invertible additive coupling layer takes as input a vector 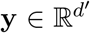, partitions it into two 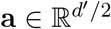 and 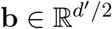, and applies

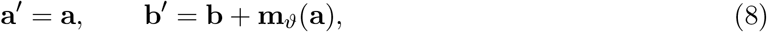

where 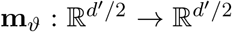 is a (possibly non-invertible) neural network. The Jacobian of this map is

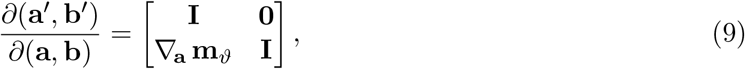

which has det = 1 identically, regardless of **m**_ϑ_. The inverse is **a** = **a**^*′*^, **b** = **b**^*′*^ − **m**_ϑ_(**a**^*′*^).

To mix the dimensions in between coupling layers, we use a Cayley-parameterized orthogonal mixer. Formally, a Cayley-parameterized orthogonal mixer takes as input a vector 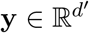 and applies an orthogonal map given by **y**^*′*^ = **Qy**, where **Q** = (**I** − **A**) (**I**+**A**) ^*−*1^ and **A** is a learnable skew-symmetric matrix. The determinant of **Q** is 1.

Thus the encoder as a whole is volume preserving (its determinant is 1), meaning it cannot arbitrarily reduce the geometric volume of the observed data to minimize noise. To handle multiple subsystems, we instantiate a separate encoder for each subsystem, so that the total encoder can be written as 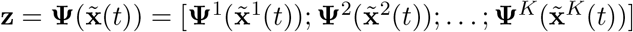.

A diffeomorphic function by construction must map to a space of the same dimensionality. However the intrinsic dimensionality of a system is in practice typically much smaller than the dimension of the delay embedding. We partition the latent space into two components: a dynamic latent 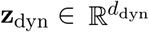 and an effectively-null latent 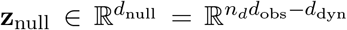, with 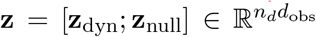 (Figure 2C, center). The null subspace is zero-padded for all decoder calls (i.e., inverse encoder calls). Thus while the null subspace is allowed to freely vary, all information for both dynamics and reconstruction must live in the dynamic latent. The JacobianODE model then operates on **z**_dyn_ only. Note that since the encoder is volume-preserving, as mentioned, all information in the input must be preserved *somewhere* in the latent space. Thus the null subspace provides the opportunity to dump any noise or information not relevant to the dynamics or reconstruction. In practice, to pick the dimensionality of the dynamic subspace *d*_dyn_, we perform a linear PCA on the delay-embedded training data and select the number of components that explain 99% of the variance. This is because linear dimensionality provides an upper bound on the intrinsic dimensionality of the system.

#### Reconstruction loss

The reconstruction loss ℒ_rec_(*θ*_Ψ_; **x**) penalizes the deviation between true and decoded latent states, where *θ*_Ψ_ are the parameters of the encoder (Figure 2C, right). We define a projection operator *P*_dyn_ that projects the latent into the dynamic subspace. Then, the reconstruction loss is given by

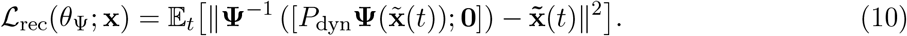

#### Latent prediction loss

The latent prediction loss ℒ_pred,**z**_(*θ*; **x**) penalizes the deviation between true and predicted latent dynamics, where *θ* = [*θ*_Ψ_; *θ*_J_] are the parameters of the encoder and JacobianODE model. Formally,

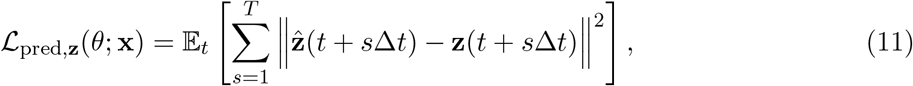

where 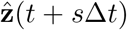 is the predicted latent state at time *t* + *s*Δ*t* and *T* is the number of prediction steps.

#### Decoded prediction loss

The decoded prediction loss _pred,**x**_(*θ*; **x**) penalizes the deviation between true and predicted observed states. To minimize the amount of redundant information, we compute the decoded prediction loss only on the most recent slice of each predicted delay-embedded window (Figure 2C, right). Formally,

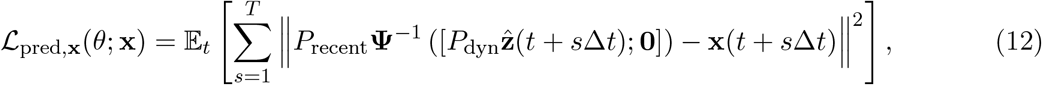

where *P*_recent_ is a projection operator that projects the decoded latent into the most recent slice of the delay-embedded window, and **x**(*t* + *s*Δ*t*) is the true observed state at time *t* + *s*Δ*t*.

#### Training objective

The total loss is given by

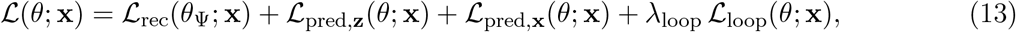

where *λ*_loop_ is a tunable hyperparameter controlling the weight of the loop-closure regularizer. Loop-closure loss is computed in latent space. The reconstruction loss and both prediction losses are given the same weight since the encoder is volume-preserving, and the relevant states should have approximately the same scale.

### 2.4 Conditional training

In the case where the data span multiple conditions, either discrete (e.g., awake vs. anesthesia) or continuous (e.g., different drug doses), we can train the JacobianODE model to be condition-aware. Specifically, we can add a conditional vector 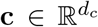 to the input of both the encoder and the JacobianODE model, and train the model to predict the Jacobian of the dynamics conditioned on this variable. In practice, we take *c* ∈ {−1, +1} to represent the difference between conditions.

## 3 Results

### 3.1 Validation on synthetic systems with ground-truth dynamics

We validated the pipeline on two simulated systems. In these systems, the true Jacobians are known, so we can compare against ground-truth invariants such as the Lyapunov spectrum, attractor topology, and ground-truth reachability and controllability gramians. The systems we used were the 3-D Lorenz attractor,^103^ and a 128-D recurrent neural network (RNN) trained on a working memory selection task inspired by Panichello and Buschman [104]. For the task-trained RNN, on each trial, the network receives two cue inputs and, after a delay and a selection cue, must sustain activation corresponding to the chosen color. The RNN consists of two 64-neuron areas (visual input and cognitive output), with predominantly within-area connectivity to encourage multi-area structure. Inputs reach only the visual area, outputs are read from the cognitive area, and thus the two areas are forced to interact to solve the task. We trained Jacobians on activity from the autonomous delay epoch before the network provides a response.^87^

Each system was trained in two variants: fully observed (the JacobianODE machinery receives all *d* state coordinates) and partially observed (the latent-space pipeline receives a *d*_obs_ ≪ *d* projection and recovers a coordinate system via delay embedding and the encoder). For the Lorenz system, only the *x* variable was observed. For the task-trained RNN, 16 coordinates were randomly chosen from each of the two areas. All models were trained on data with 5% observation noise.

We evaluated trajectory prediction via *R*^2^ and decoder-corrected mean absolute standardized error (MASE). The decoder-corrected MASE normalizes the mean absolute error by the absolute error of the decoded persistence baseline 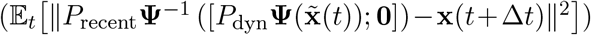, which separates the dynamic prediction quality from the decoder error. Trajectory-prediction quality is reported on held out test data (Figure 3A). In both the fully-observed and partially-observed cases, the models are able to reproduce trajectories, though this ability is predictably somewhat impaired for the partially-observed Lorenz system.

**Figure 3.**
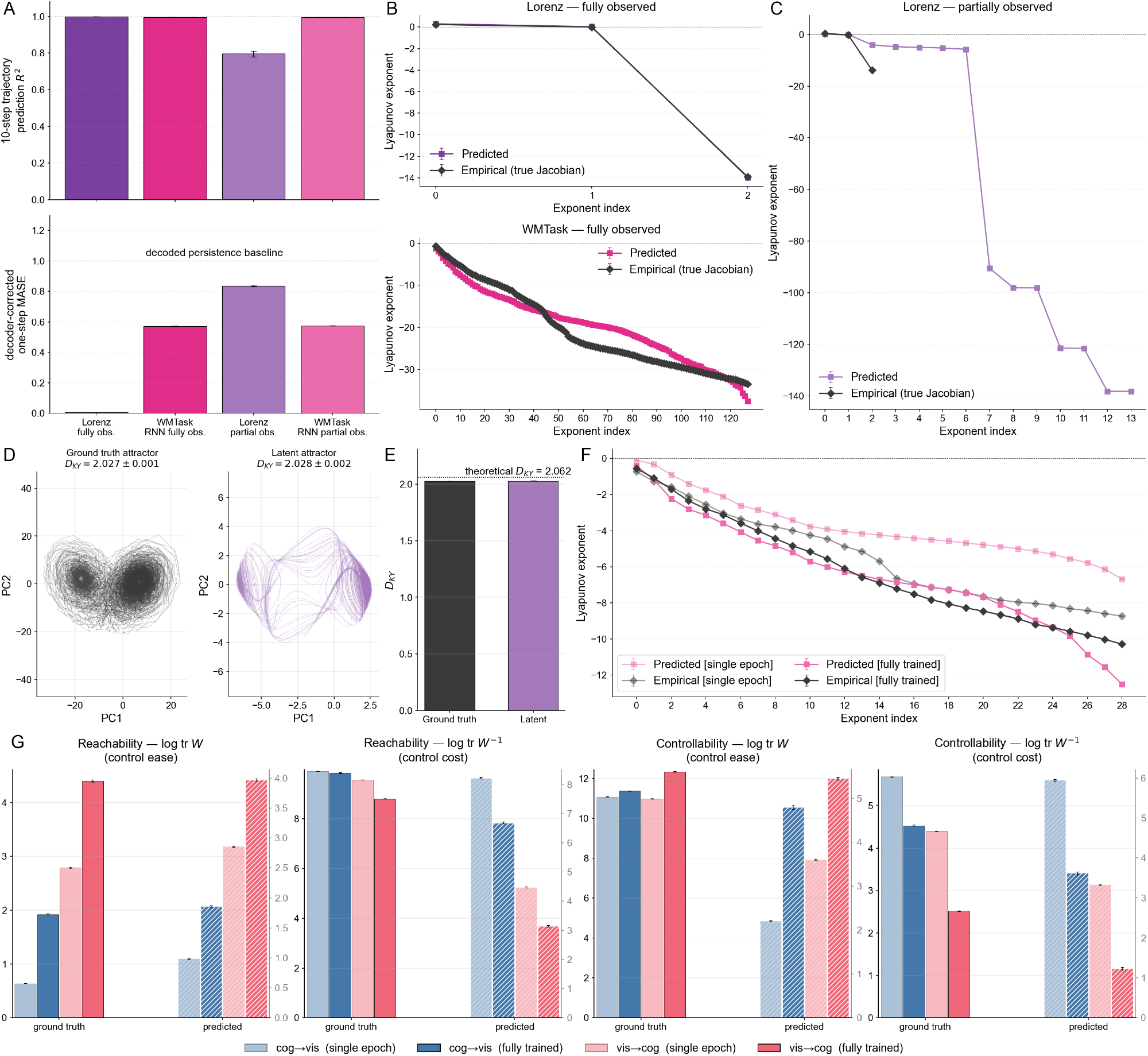
Validation on synthetic systems with ground-truth dynamics. (A) Ten-step free-running *R*^2^ (top) and decoder-corrected one-step MASE (bottom) for the four runs. (B) Predicted vs. empirical Lyapunov spectra (sorted descending) for the two fully observed runs. (C) Same for Lorenz partially observed. (D–E) Ground-truth and latent-recovered Lorenz attractor in PC1–PC2 with Kaplan– Yorke dimension annotations. (F) Task-trained RNN single-epoch versus fully trained Lyapunov spectra, predicted (pink) vs. empirical (black). (G) Task-trained RNN partially observed directional reachability and controllability log-trace and inverse log-trace, ground-truth vs. predicted, per condition.

The Lyapunov spectrums are recovered by the latent JacobianODE models in the fully observed case (Figure 3B). For the partially-observed Lorenz system, the latent JacobianODE formulation recovers the top two exponents, while distributing the third exponent, corresponding to decay onto the attractor, into the remaining 12 latent dimensions (Figure 3C). Note that the partially observed latent spectrum has 14 entries due to the method of picking the latent dimension *d*_dyn_ by PCA on the delay-embedded training data at variance threshold 0.99. The results reflect that under partial observation, the encoded delay-embedded states do not contain the necessary resolution to resolve the third exponent exactly. However, the latent JacobianODE does recover the Kaplan-Yorke dimension of the Lorenz attractor, indicating that the attractor topology is preserved (Figure 3D,E). For the partially observed task-trained RNN, the latent JacobianODE model was trained conditioned on the condition variable *c* ∈ {− 1, +1}, indicating the first epoch or the final epoch of training. The latent JacobianODE model qualitatively recovers the leading exponents of the Lyapunov spectrum in each condition, as well as the relevant difference between them (Figure 3F).

The latent JacobianODE model also recovers the relative changes in reachability and controllability between the two conditions (Figure 3G). Reachability increases from the first epoch to the final epoch of training for both visual to cognitive control and cognitive to visual control (Figure 3G, left and center-left). Additionally, visual to cognitive reachability is higher in both conditions, reflecting the need for the visual area to drive the cognitive area to make the correct decision. Controllability also increases from the first epoch to the final epoch of training for both visual to cognitive control and cognitive to visual control (Figure 3G, center-right and right).

### 3.2 JacobianODE models capture neural dynamics and show more nonlinear geometry in awake state relative to anesthesia

We applied the pipeline to multi-area LFP from two non-human primates (NHPs) recorded during propofol infusion. Each session simultaneously recorded LFP from posterior parietal cortex (PPC), superior temporal gyrus (STG, auditory cortex), frontal eye fields (FEF), and ventrolateral pre-frontal cortex (vlPFC) at 1 kHz. We low-pass filtered the signals at 80 Hz. We used resting state trajectories for training the JacobianODE model, defined as starting at least 3 seconds after the onset of any external stimulus (either airpuff, or auditory tone). Data was taken from either the awake state or a maintenance dose anesthetic state, and the JacobianODE model was conditioned on the condition variable *c* ∈ {−1, +1}, indicating the state.

Free-running trajectory prediction and decoder-corrected one-step MASE are reported per condition, per NHP (Figure 4A). Trajectories are well captured over 10-step free-running prediction. Anesthetic state trajectories were generally easier to predict than awake state (*p* = 9.5 × 10^−7^, paired two-sided Wilcoxon signed-rank test, median difference anesthesia − awake Δ*R*^2^ = 0.068 [95% CI 0.060, 0.071]), though for both conditions the decoder-corrected one-step MASEs are below the decoder-corrected persistence baseline of 1.0.

**Figure 4.**
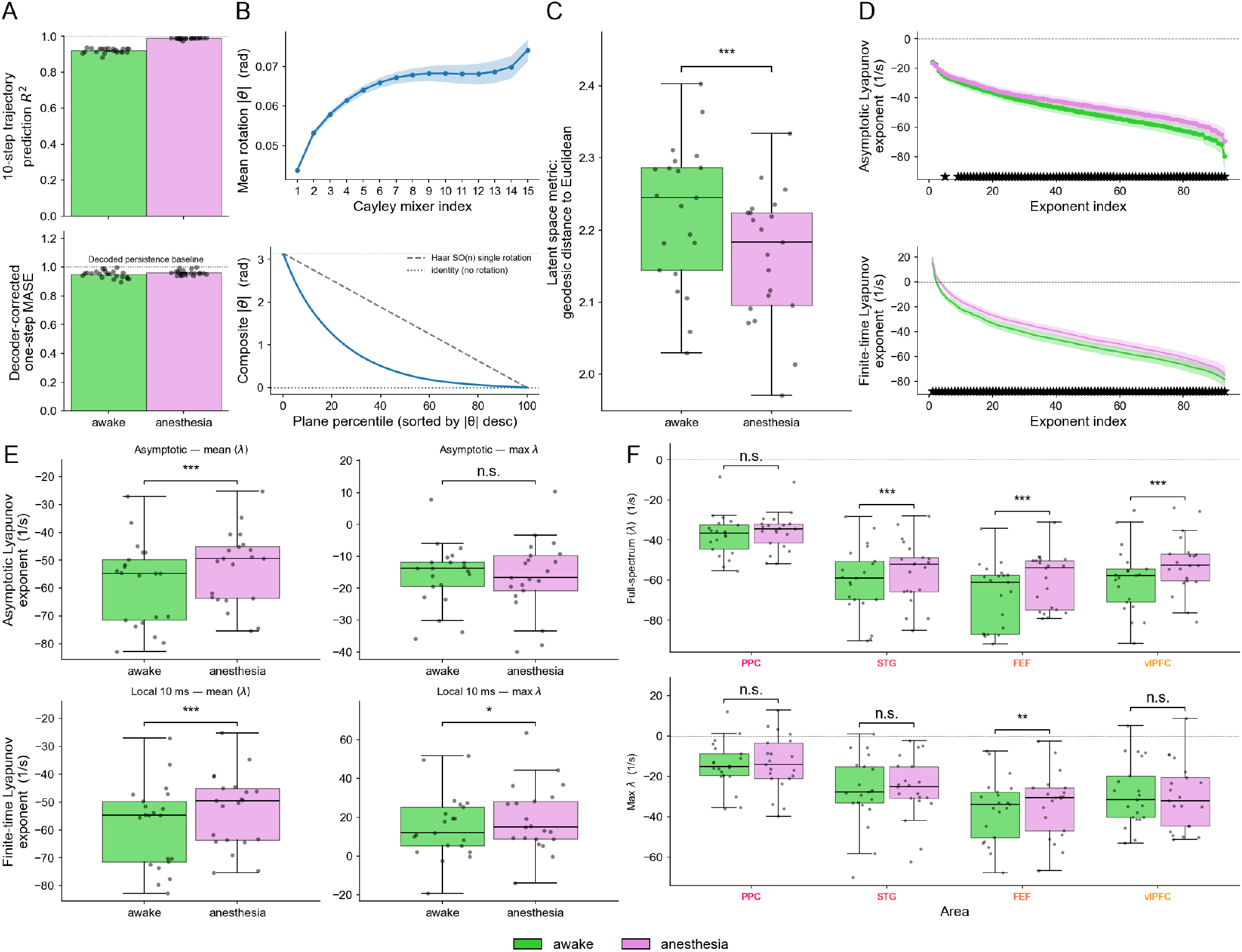
Trajectory prediction, encoder geometry, and Lyapunov spectra under propofol (*n* = 21 sessions). (A) Per-session 10-step *R*^2^ and decoder-corrected one-step MASE, awake vs. anesthesia. (B) Mean per-Cayley-layer rotation angle through encoder depth, between identity (*θ* = 0) and the Haar SO(*n*) reference. (C) Geodesic distance from identity of the composite encoder Jacobian per session, awake (larger; more warped metric) vs. anesthesia (*p <* 0.001). (D) Full asymptotic (top) and 10-ms FTLE (bottom) Lyapunov spectra per session, sorted descending. (E) Boxplots of asymptotic mean and max exponents and of 10-ms FTLE mean and max exponents. (F) Per-area within-area asymptotic mean (top) and max (bottom) exponents per session for PPC, STG, FEF, vlPFC; sign-agree-gated significance annotated.

The encoder leverages considerable transformations to encode the dynamics into the latent space. The angle of rotation rises through the depth of the encoder, and rotates the latent space into a more complex geometry (Figure 4B). Furthermore, the Riemannian metric induced on the latent space is farther from identity in the awake state than during anesthesia, indicating a more complex geometry in the awake state (Figure 4C; *p* = 9.5 × 10^−7^, paired two-sided Wilcoxon signed-rank test, median difference anesthesia awake Δ = − 0.059 [95% CI− 0.067, − 0.050]). This suggests that the decreased prediction quality in the awake state relative to anesthesia may be related to the more nonlinear geometry of the data manifold.

### 3.3 Broad-spectrum destabilization during propofol anesthesia

Previous work has shown that neural dynamics are destabilized under propofol anesthesia.^92,93^ We confirmed this destabilization using the latent JacobianODE models. We computed Lyapunov exponents for all resting state trajectories within each epoch of each session and then averaged across sessions. We report both the asymptotic Lyapunov spectrum as well as the finite time Lyapunov exponent (FTLE) spectrum, which was computed over 10 millisecond windows.

While the largest asymptotic Lyapunov exponents are similar across conditions, the broad spectrum of asymptotic Lyapunov exponents was significantly destabilized under anesthesia, as was previously found (Figure 4D,E; *p* = 9.5 × 10^−7^, paired two-sided Wilcoxon signed-rank test, median difference full spectrum mean anesthesia − awake Δ ⟨*λ*⟩ = +5.61 [95% CI +4.56, +6.78]). For the finite-time Lyapunov exponents, both the maximum Lyapunov exponent and full spectra show destabilizations during anesthesia (Figure 4D,E; max exponent *p* = 0.038, paired two-sided Wilcoxon signed-rank test, median paired difference, anesthesia awakeΔ⟨*λ*⟩ = +3.82 [95% CI +0.52, +6.97], full spectrum *p* = 9.5 × 10^−7^, ⟨*λ*⟩ Δ = +5.61 [95% CI +4.56, +6.78]). We note that the maximum Lyapunov exponents are not always faithfully reproduced in the sample systems and may be more sensitive to noise. They should therefore be interpreted with caution. The full spectrum however was robustly recovered by JacobianODE models, and more consistently depicts the destabilization across a multitude of directions.

Per-area asymptotic Lyapunov exponents qualitatively reproduce the shift found at the full system level (Figure 4F). Notably, PPC had the most unstable dynamics both in the awake and anesthetic states (*p* ≤ 3×10^−6^, paired two-sided Wilcoxon signed-rank test).

### 3.4 Propofol anesthesia reduces directional reachability and controllability

We next sought to characterize how propofol anesthesia changes interareal reachability and controllability. We computed the directional reachability and controllability Gramians for each condition using the learned Jacobians. We report both the reachability and controllability Gramians computed over 10 millisecond horizons. Each gramian measures the influence of area *β* on area *α* through the time-dependent coupling, defined by **J**^β*→*α^. For 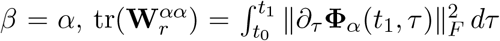 and 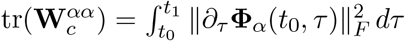: the integrated squared Frobenius norms of how the within-area state-transition matrices evolve with the integration time. This can be interpreted roughly as the strength of the within-area recurrence, and provides a natural baseline for comparing the trace magnitudes.

Reachability was broadly decreased under anesthesia, both when considering the easiest directions of control (Figure 5A, left) and the most difficult directions of control (Figure 5A, right). Notably, the reachability along the most manipulable directions became *easier* under anesthesia for the interactions involving both PPC and FEF driving vlPFC. Controllability was also broadly decreased under anesthesia, both when considering the easiest directions of control (Figure 5B, left) and the most difficult directions of control (Figure 5B, right).

**Figure 5.**
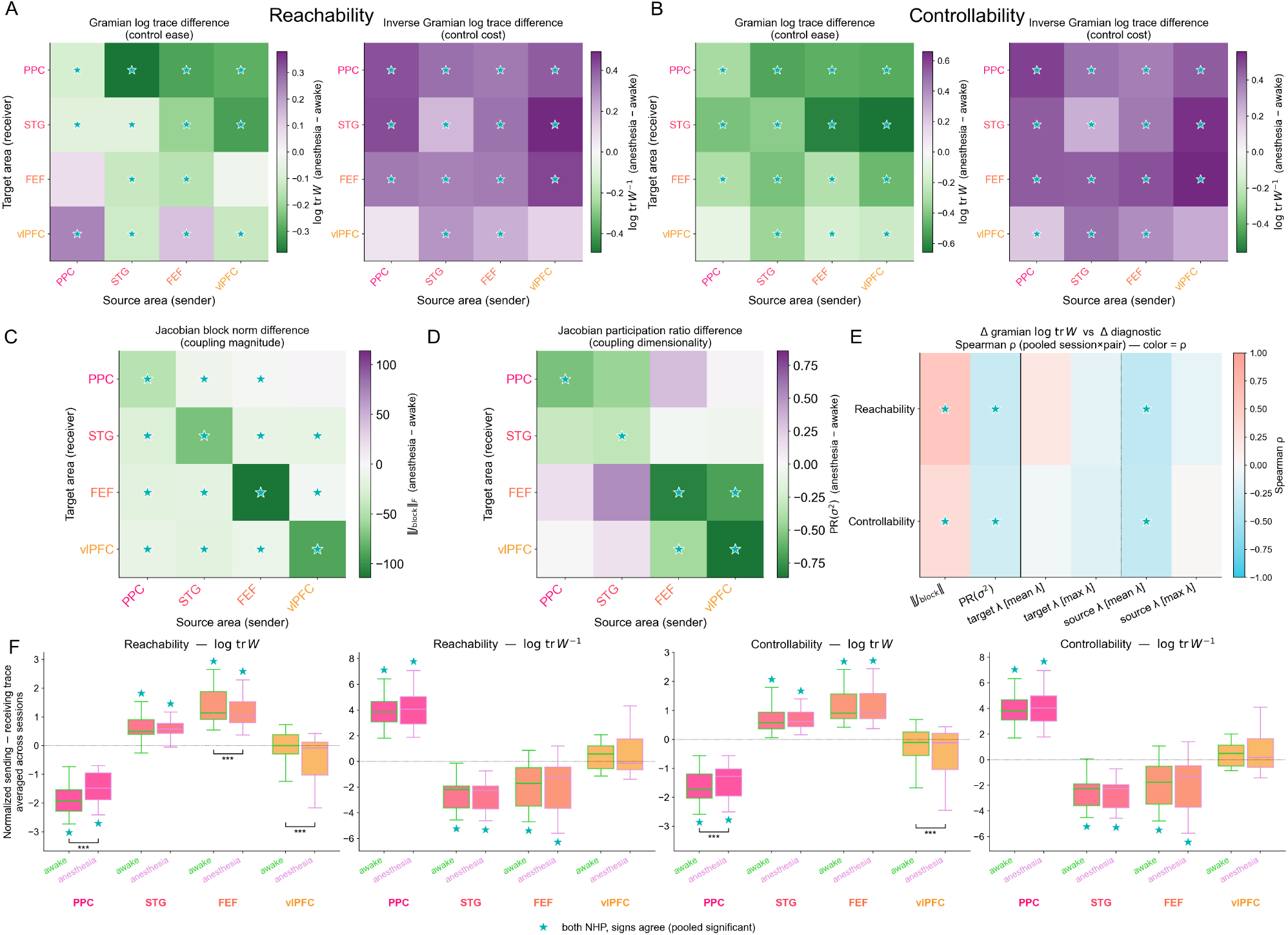
Directional reorganization of interareal control under propofol (*n* = 21 sessions). (A) 4 × 4 source–target heatmaps of pooled anesthesia − awake differences in reachability log tr **W** (left) and log tr **W**^*−*1^ (right); stars mark cells passing the sign-agree + BH-FDR gate at *q <* 0.05. (B) Same for controllability. (C) Jacobian block Frobenius-norm differences. (D) Block participation-ratio differences. (E) Spearman correlation between Gramian-change cells and block-norm/PR-change cells. (F) Per-area net send − receive axis normalized by diagonal within-area measure across all four metrics, placing PPC at the receiver end, FEF/STG at the sender end, vlPFC balanced.

These disruptions in reachability and controllability are most evident at the short time horizon (10 milliseconds) presented here. At longer horizons, the intrinsic dynamics (more destabilized under anesthesia) increasingly dominate the Gramians over the interareal coupling. When the target system is less stable, reachability becomes easier and controllability becomes harder. Thus, over longer horizons, reachability becomes easier under anesthesia (particularly for projections to PPC and STG), while the controllability reduction instead deepens (Appendix A.2). This illustrates that a reduction in the ability to drive posterior areas manifests on shorter horizons (10, 20, and 100 milliseconds), while on longer timescales (1 second) the intrinsic instability of the target area dynamics leads to greater reachability.

### 3.5 Anesthesia reduces coupling strength and local coupling dimensionality

To probe the influence of anesthesia on coupling strength, we analyzed how anesthesia changes both the Frobenius norm of the cross-area and within-area Jacobian blocks (Δ log tr **W** and Δ ∥ **J**_block_ ∥, the difference in Frobenius norm of the coupling blocks between states). We also analyzed the participation ratios of these blocks (ΔPR(*σ*^2^), i.e. change in the participation ratio of the singular values of 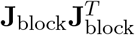). We found that the Frobenius norm of the Jacobian blocks broadly decreased during anesthesia, indicative of reduced coupling strength between areas (Figure 5C). The participation ratios (dimensionality) of the Jacobian blocks showed more mixed results. Within frontal (FEF, vlPFC) and within posterior (PPC, STG) cortices anesthesia decreased coupling dimensionality (Figure 5D). All four within-frontal pairs decreased significantly. For the frontal-to-posterior and posterior-to-frontal couplings, the results were not significant but tended to be higher during anesthesia. This suggests that anesthesia decouples more closely related areas but potentially creates more diffuse, albeit weakened, couplings across more distant areas. Indeed, within-cluster coupling was weakened more by anesthesia than between-cluster coupling, (*p* = 9.5×10^−6^, paired two-sided Wilcoxon signed-rank test, median difference within-cluster − between-cluster Δ[Δ*PR*(*σ*^2^)] = −0.69 [95% CI −0.94, −0.43]).

### 3.6 Changes in reachability and controllability are driven by multi-factorial anesthesia-induced effects

We assessed how various factors explain changes in reachability and controllability. We correlated the changes in Gramian traces with changes in block-norm and participation-ratio, as well as with changes in target and source-area Lyapunov exponents across conditions (Figure 5E). The traces are positively correlated with block-norm changes, indicating that coupling magnitude is a significant contributor to control ease. Interestingly, they are negatively correlated with the participation ratio changes, indicating that more diffuse coupling might actually impede control along the most essential directions. Overall, the correlations were relatively weak, reflecting that directional control depends on a multitude of factors that include the coupling strength and dimensionality, as well as the dynamic stability of the source and target areas, and the alignment of the source projection with the directions of flow in the target area.

### 3.7 Anesthesia disrupts the hierarchy of cortical organization into senders and receivers

Finally, we evaluated whether anesthesia changes area-level patterns of sending and receiving. We first normalized each cross-area gramian measure by the within-area measure of the target area. We then computed the column-wise mean (corresponding to average sending capability) and row-wise mean (corresponding to average receiving capability). This gives us a per-area measure of the average sending capability relative to receiving capability within each condition (Figure 5F). We found that across both awake and anesthesia, PPC was consistently more of a receiving area, while STG and FEF were consistently more sending areas. vlPFC was balanced, indicative of its role as a hub. While the roles were preserved across conditions, anesthesia tended to push areas more toward a balanced regime (sending capability equivalent to receiving capability), thus disrupting cortical organization. PPC and FEF, the two areas with the most extreme awake-state roles, both shifted toward balance under anesthesia.

## 4 Discussion

Our results show that anesthesia decreases the ability of areas to control each other, both in terms of driving towards novel states and stabilizing along existing trajectories. We reproduced earlier results, showing that propofol anesthesia induces broad spectrum destabilization of neural dynamics. While the destabilization was most pronounced in PPC, it was present across all areas. Interestingly, despite decreased stability correlating with increased reachability ease in the absence of other changes, we found that reachability broadly decreased in anesthesia. This suggests that at the time horizon of 10 milliseconds considered in the main text, the decreased stability in target areas could not compensate for the reduction in coupling strength also observed. This again highlights that changes in reachability and controllability were driven by multi-factorial anesthesia-induced effects, including changes in coupling strength and dimensionality, as well as the dynamic stability of the source and target areas.

### Paradoxical excitation

Notably, the directional interactions from PPC to vlPFC and from FEF to vlPFC showed surprising increases in reachability under anesthesia. These pathways could potentially be interpreted as a mechanism of the paradoxical excitation observed during propofol anesthesia. Paradoxical excitation is characterized by an increase in abnormal movements as well as sensory hallucinations.^105–107^ Thus, it may be that the increased ease with which PPC and FEF can drive vlPFC under anesthesia may lead to the observed spurious movements and sensory hallucinations prior to the onset of loss of consciousness.

### Reduced coupling magnitude during anesthesia

Our finding of reduced coupling magnitude during anesthesia aligns with previous findings of reduced synchrony and network integration during propofol infusion.^37,45,49–64^ Furthermore, the finding that coupling dimensionality decreases locally (within frontal and posterior areas) but increases more globally (across frontal and posterior areas) aligns with recent findings of reduced synchrony within hemisphere but increased synchrony across hemispheres.^49^

### Neuropsychiatric conditions

We also note that the latent JacobianODE pipeline leveraged in this work was agnostic to both the recording modality and the nature of the data (anesthetic infusion), and could be applied in a wide range of settings. Neuropsychiatric conditions, in particular, present a compelling setting in which to explore the application of nonlinear interareal control analyses. Neural control has been shown to be relevant in schizophrenia,^77,84,108^ and epilepsy,^78^, and a lack of goal-directed control has been identified across multiple psychiatric conditions.^109^ Additionally, dysregulation of the cortico-basal ganglia-thalamocortical loop has been associated with a wide range of neuropsychiatric conditions that includes addiction, attention-deficit/hyperactivity disorder, Parkinson’s, and many others.^110–112^ Notably, these conditions all involve a lack of control in some form, suggesting this might be a compelling setting in which to look for control theoretic biomarkers and treatment predictors.

### Interventional control

Prior work has shown that JacobianODE models can be used to directly control synthetic neural models to produce desired behaviors.^87^ This raises the possibility of using the latent JacobianODE pipeline for interventional control in both basic science and clinical settings, such as transcranial magnetic stimulation (TMS) and deep brain stimulation (DBS).^113–117^ Given an appropriate mapping from stimulus to brain activity, the control inputs could actually be provided as sensory stimuli, obviating the need for technologically complex manipulation.^118^

### 4.1 Future directions

We validated that the latent JacobianODE method is capable of capturing relative changes in reachability and controllability from partial observation in the task-trained RNN. However, it should be determined whether the model is capable of recapitulating these relative changes when there are more than two subsystems present.

A challenge of the latent Jacobian estimation pipeline is the need to pick the intrinsic dimensionality of the latent dynamics *d*_dyn_. In the present work, we err on the side of caution and pick a dimensionality sufficient to explain 99% of the variance in principal component space in the data. However a nonlinear method of intrinsic dimensionality estimation that yields tighter bounds to the true intrinsic dimensionality of the system would both save compute (as higher dimensions require larger models) and likely yield improved Jacobian estimation. An outstanding possibility not explored in this work is to use the learned geometry of the encoder (the induced Riemannian metric on the latent space) to infer the true number of directions that are relevant for the latent manifold.

Furthermore, the models fit to neural data surpassed the decoder-corrected persistence baseline, but did not adequately reproduce the asymptotic dynamics of the system. To do so, a compelling alternative is to leverage the nonautonomous extension of the JacobianODE framework suggested in the initial presentation of the method.^87^ Incorporating this change would vastly increase the expressivity of the model and potentially lead to more effective models of neural dynamics.

While we focus here on control, we note that a wide range of tools have been developed and harnessed to study interareal interactions in neural data.^119,120^ They include methods based on reduced rank regression,^121,122^ recurrent neural network models of neural dynamics,^123–127^ Gaussian-process factor analysis,^128^ canonical correlation analysis,^129,130^ convergent cross mapping,^131,132^ switching dynamical systems,^133,134^ granger causality,^135,136^ and point process models.^137^ A complete analysis should consider how these existing metrics vary with propofol anesthesia to provide a more comprehensive picture of changes in interareal interactions during anesthetic-induced unconsciousness.

## Methods: Disruption of interareal control during propofol anesthesia

### A Supplementary results

#### A.1 Linearity of the learned dynamics and linear baseline analysis

A natural question raised by the conditional JacobianODE model is whether the dynamics it learns from the neural data are genuinely nonlinear, or whether a strictly linear time-invariant (LTI) description of the same data would suffice. We address this question in two ways (Figure S1). We first investigate how strongly the learned Jacobian vary along the trajectory, and how that variation trades off against predictive accuracy (Figure S1A,B). Second, we retrain the dynamics with a linear delay-embedded model and ask whether the same anesthesia-related Gramian reorganization is retained (Figure S1C,D).

##### The learned Jacobian is approximately linear, despite targeted anti-LTI interventions

We compared the baseline JacobianODE model against five variants designed to push the Jacobian off the constant-matrix solution: separate per-condition dynamics, a per-batch latent exponential moving average normalization, a null-space penalty, a stop-gradient applied to the loop-closure target, and a smaller encoder. As a non-LTI reference we include the partially observed workingmemory selection RNN of §D.1.2 (a 128-D task-trained network whose ground-truth dynamics are genuinely nonlinear). For each model we report three diagnostics on the validation set: the per-condition Jacobian temporal coefficient of variation, 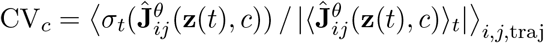, the loop-closure loss ℒ _loop_, and the free-running trajectory validation loss (Figure S1A). The baseline model had a relatively low temporal CV, indicating approximately LTI latent dynamics. Across all five interventions, temporal CV stays within a factor of two of the baseline (CV ≈ 10^−3.5^–10^−3^), three orders of magnitude below the task-trained RNN reference (CV ≈ 3). Thus the variation of **Ĵ**^θ^ along the trajectory is small even when the architecture is explicitly biased against the constant-matrix solution. None of the interventions yields a meaningful reduction in ℒ _loop_ or trajectory loss either. The model is therefore capable of representing strongly nonlinear Jacobians when the data demand it (the task-trained RNN reference confirms this), but on the propofol cohort it settles into the approximately-LTI regime. This may have to do with the construction of the dynamics as autonomous, or with the short prediction horizon (10 milliseconds) used for training.

##### The overall observation-space dynamics remain nonlinear through the encoder and decoder

The near-LTI behaviour of **Ĵ**^θ^ is a statement about the *latent* dynamics alone. The dynamics in the observed delay-embedded space need not be linear, because they are read out through the nonlinear diffeomorphic encoder. The model advances a delay-embedded observation by encoding it, evolving the latent state by encoding, integrating **Ĵ**^θ^, then decoding 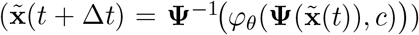. By the chain rule the observation-space Jacobian therefore factorizes as

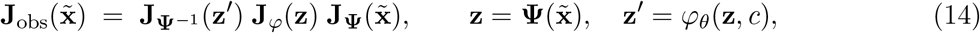

which is the product of the encoder Jacobian, the latent flow Jacobian, and the decoder Jacobian. Even when the central factor **J**_φ_ is nearly state-independent, the encoder and decoder Jacobians **J**_**Ψ**_ and 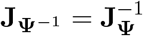 vary with the state, so the composite observation-space map is genuinely nonlinear. To quantify this we computed the same per-condition temporal CV for the encoder Jacobian and the (exactly invertible) decoder Jacobian on the held-out validation set (Figure S1A, top row; for NHP 1 the decoder Jacobian is obtained as 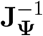, which matches a direct automatic-differentiation Jacobian of **Ψ**^*−*1^ to a relative error of ∼ 10^−14^). For NHP 1 the encoder and decoder Jacobians vary substantially along the trajectory. The near-constancy of **Ĵ**^θ^ thus reflects the encoder absorbing the nonlinearity of the observed dynamics into a coordinate system in which the latent flow is approximately linear, not the observed neural dynamics being linear.

This view connects the latent JacobianODE to Koopman operator theory, in which a nonlinear system is rendered linear by lifting it through a map of observables^138,139^, and in particular to deep-learning approaches that learn such linearizing coordinates with autoencoders^140,141^. The diffeomorphic encoder differs in being an exactly invertible, volume-preserving change of coordinates of the same dimension, rather than a (generally higher- or infinite-dimensional) lift to a space of observables; correspondingly the latent flow is only *approximately* linear, since an invertible, dimension-preserving change of coordinates cannot in general render a nonlinear flow exactly linear (for instance, when it possesses multiple isolated fixed points).

##### Loop-closure regularization trades nonlinearity against predictive accuracy

We found that as the loop closure loss weight *λ*_loop_ was varied, the best free-running trajectory validation loss was monotonically non-decreasing for both NHPs. Increasing *λ*_loop_ enforces row-conservativity of **Ĵ**^θ^, an integrability constraint that the constant-matrix solution satisfies trivially. Together with panel A, this is consistent with the interpretation that, for the propofol data, a near-LTI Jacobian is the most parsimonious model: gains in nonlinearity (lower ℒ _loop_) come at the cost of a worse fit.

##### Linear delay-embedded baseline: per-area eigen-time-delay coordinates with joint DMD

Given the near-LTI behaviour of the JacobianODE model on the cohort, we asked whether substituting a strictly linear delay-embedded model for the learned Jacobian preserves the per-pair Gramian readouts reported in the main text. The baseline is a per-area extension of the Hankel Alternative View Of Koopman (HAVOK) construction^142,143^, fit as follows. For each area *k* ∈ {1, …, *K*} and each time window (§D.3), we form the per-area Hankel matrix at depth 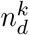 from the 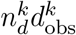 stacked delay coordinates of §C.1.1, stack the windows of one condition, and take its reduced SVD to obtain the per-area eigen-time-delay basis:

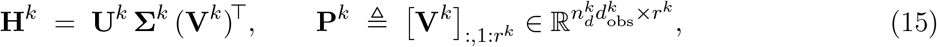

so that the per-area reduced state is 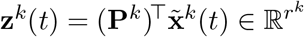. The pair (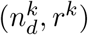) is selected per area by an AIC sweep over 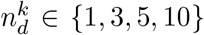 and *r*^k^ ∈ {4, 8, 16, …, 600} on held-out within-condition windows (80*/*20 window-level split, per-area observation-space prediction error). Reduced states from the *K* areas are concatenated to a joint state **z**(*t*) = [**z**^1^(*t*); … ; **z**^K^(*t*)] ∈ ℝ ^R^ with *R* = _k_ *r*^k^ (aligning all windows to the maximum 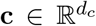 leading offset so the per-area snapshot indices align), and a joint discrete-time linear model is fit by least-squares dynamic mode decomposition,

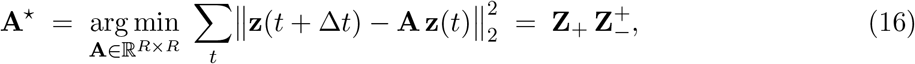

where **Z**_*−*_ = [**z**(*t*_1_), …, **z**(*t*_N*−*1_)] and **Z**_+_ = [**z**(*t*_2_), …, **z**(*t*_N_)] are the stacked snapshot pairs and (·)^+^ denotes the Moore–Penrose pseudoinverse. The construction of the *K*-area joint state imposes a natural block decomposition of **A**^⋆^: writing 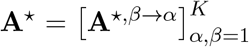 with 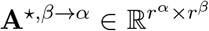 acting as the linear analogue of **J**^β*→*α^ used in the main text, the discrete-time reachability and controllability Gramians at horizon *T* are computed by the square-root QR iteration of §C.4.2 with (**A**_k_, **B**_k_, **C**_k_) = (**A**^⋆,α*→*α^, **A**^⋆,β*→*α^, **A**^⋆,α*→*β^) time-invariant (the same numerical pipeline, with the constant LTV reference replaced by the fitted **A**^⋆^). All Gramians reported below use horizon *T* = 10Δ*t*, matching the main text’s JacobianODE horizon.

##### The linear baseline reproduces the JacobianODE ordering on the task-trained RNN

Applied to the partially observed task-trained working-memory selection RNN of §D.1.2, the per-area linear HAVOK pipeline qualitatively reproduces the ground-truth pattern of reach and controllability Gramian magnitudes for the → visual cognitive and cognitive → visual blocks, both at the single-epoch and fully-trained checkpoints used in the main text (Figure S1C). While the directional ordering between conditions is not perfectly preserved on every panel, the linear baseline is informative on a nonlinear test bed, and is therefore an appropriate sanity check to run on the neural data.

##### The linear baseline reproduces the main propofol findings, with reduced precision

We ran the same per-area linear pipeline across all 21 sessions of the propofol cohort and computed the four sign-agree pooled-Wilcoxon log tr **W** and log tr **W**^*−*1^ heatmaps that constitute Figure 5A,B of the main text (Figure S1D, 10 ms integration horizon). The qualitative organization survives: anesthesia decreases reachability and controllability ease (more negative Δ log tr **W**) and increases their associated cost (Δ log tr **W**^*−*1^) on most pairs, with the same notable exception that anesthesia *increases* reach ease from PPC and FEF into vlPFC. Compared to the JacobianODE composite, however, the linear baseline shows more variability and smaller effect sizes. This is the expected signature of a less precise model: a single time-invariant block matrix is a coarser representation of the data than a per-trajectory conditional Jacobian, but the directional reorganization the main text reports is robust to this substitution.

#### A.2 Interareal control reorganization across integration horizons

The directional control summaries reported in the main text use a *T* = 10 ms integration horizon (§C.4). The gramian integrand combines a cross-area input term (the coupling block **B** = **J**^β*→*α^) with the within-target state-transition propagator **Φ**^α^ (Eqs. 65–66), thus lengthening the horizon progressively up-weights the intrinsic within-area dynamics relative to the interareal coupling. To ask whether the anesthesia-driven reorganization is a property of the coupling or of the intrinsic dynamics, we recomputed the full set of summaries (reachability and controllability log tr **W** and log tr **W**^*−*1^, and their diagnostic correlations) at integration horizons of 10, 20, 100 and 1000 ms (Figure S2).

At the control-relevant 10 ms horizon, as reported in the main text, anesthesia shrinks reachability between areas (Figure S2, top row). This reachability reduction progressively dissolves as the horizon lengthens: the number of pairs with a significant reachability log tr **W** decrease falls from 12 (10 ms) to 9 (20 ms) to 7 (100 ms) to 3 (1 s), and by 1 s the pooled mean difference has reversed sign (pooled mean Δ = +0.22, vs. − 0.11 at 10 ms). Controllability behaves oppositely: the trace decrease not only persists but amplifies monotonically with horizon, as the longer integration window accumulates more of the within-target propagator relative to the bounded cross-area input.

**Figure S2.**
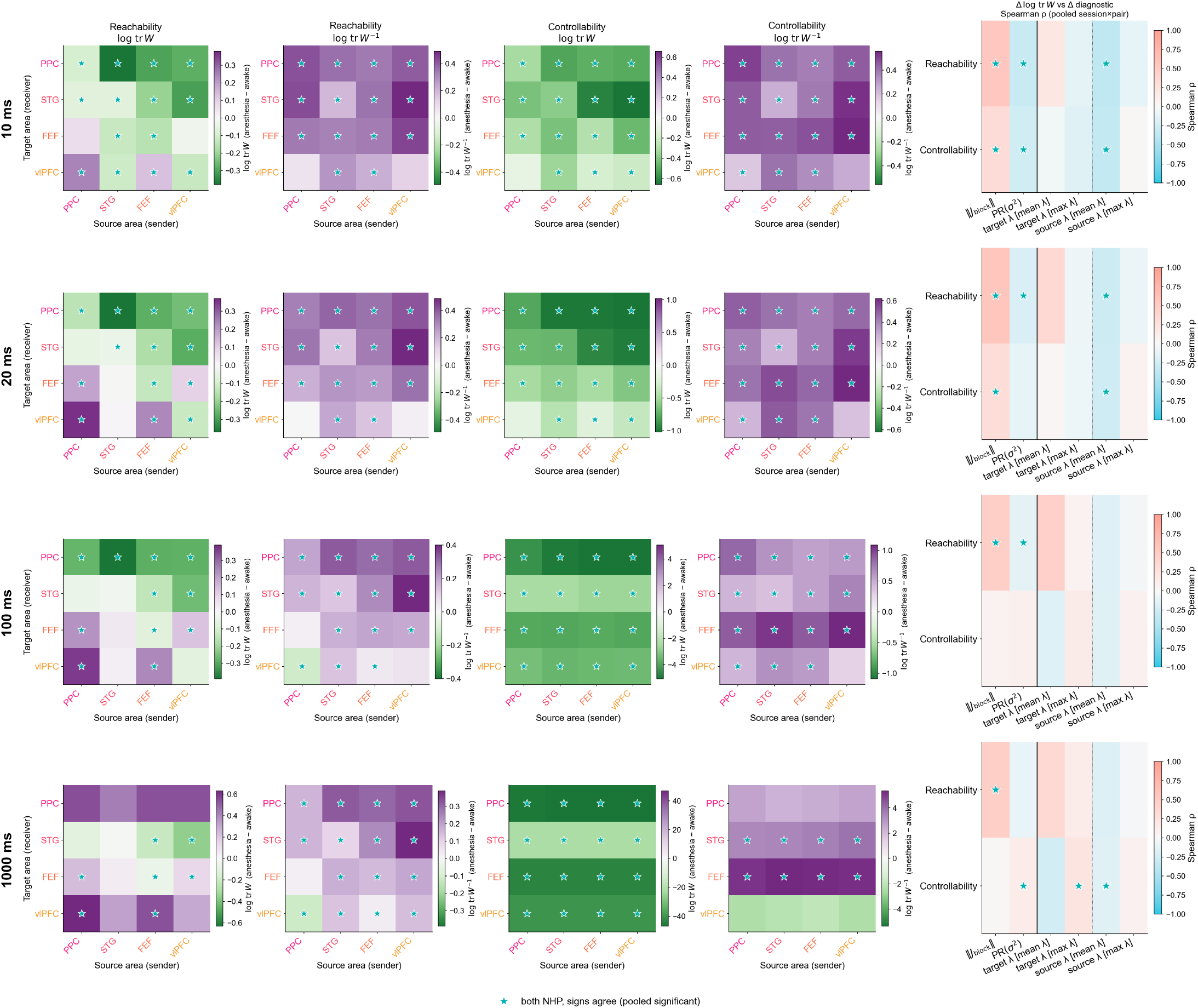
Interareal control summaries across integration horizons (*n* = 21 sessions). Rows: gramian integration horizon (10, 20, 100, 1000 ms). Columns 1–4: 4 × 4 source–target heatmaps of pooled anesthesia awake differences in reachability log tr **W** and log tr **W**^*−*1^ and controllability log tr **W** and log tr **W**^*−*1^; teal stars mark cells passing the sign-agree + BH-FDR gate at *q <* 0.05 (same convention as Fig. 5A,B). Column 5: Spearman correlation between the per-pair change in Gramian log tr **W** and the change in block-norm, participation ratio, and source- and target-area Lyapunov exponents, with session-clustered permutation testing (same construction as Fig. 5E), computed at that horizon. The 10 ms row reproduces the main-text summaries.

The diagnostic correlations tell the same story from the explanatory side. At 10–20 ms, the change in reachability tracks the change in cross-area coupling magnitude (Δ ∥**J**_block_ ∥, *ρ* = +0.56), coupling dimensionality (ΔPR(*σ*^2^), *ρ* = − 0.31), and source-area stability (Δsource-area *λ, ρ* = − 0.36). As the horizon lengthens, the source-area-*λ* and participation-ratio associations vanish, and the only correlate that remains stable across all four horizons is coupling magnitude (*ρ* = +0.56, +0.57, +0.48, +0.47). A target-area-*λ* association emerges at the longer horizons but does not reach significance (*ρ* = +0.45 at 100 ms, *p* = 0.08). The explanatory weight thus migrates from the input coupling toward the target area’s own dynamics as the integration window grows.

Together, these trends indicate that the anesthesia-driven reachability reorganization reported in the chapter is a short-horizon, coupling-governed effect. Over longer windows, the within-area state-transition propagator **Φ**^α^ ∼ exp(**J**^α*→*α^ *t*) (the intrinsic dynamics whose leading Lyapunov exponents are characterized in §G) increasingly dominates the Gramian over the cross-area input term, reshaping the apparent reachability landscape (the sign reversal at 1 s). At the same time, the destabilized anesthetic state drives the controllability ease even further down over long horizons. Crucially, the increased reachability capability under anesthesia is past the timescale on which one area can steer another before the target’s own dynamics carry its state away: by the point at which intrinsic dynamics set the reachability landscape, the window for directed interareal control has likely effectively closed. The reachability disruption (reduced reachability under anesthesia, set by the interareal coupling) is the one visible at the short, functionally relevant horizon.

### B Proofs

#### B.1 Jacobian-parameterized time derivative

The trick that makes JacobianODE estimation backpropagate cleanly through **Ĵ**^θ^ rather than through a separately-parameterized **f** is to express the instantaneous time derivative as a functional of the Jacobian alone, evaluated along a smooth path between two observed states. By the fundamental theorem of calculus along the trajectory,

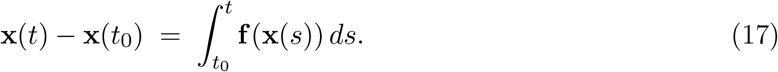

The conservativity-induced path-integral identity (Eq. 5) lets us replace each integrand **f** (**x**(*s*)) by **f** (**x**(*t*)) minus the path integral of **J** from **x**(*s*) to **x**(*t*), and solving the resulting linear relation for **f** (**x**(*t*)) gives the central proposition used throughout JacobianODE training.

##### Proposition 1

(Jacobian-parameterized **f**). *Given observed states* **x**(*t*_0_) *and* **x**(*t*) *along a trajectory of* 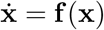 *with continuous Jacobian* **J** = ∇**f**, *there exists a functional* F *of* **J** *alone such that*

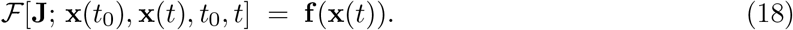

*Explicitly*,

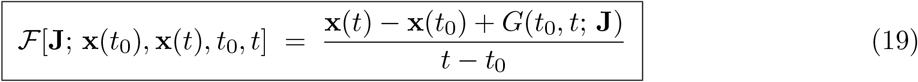

*Where*

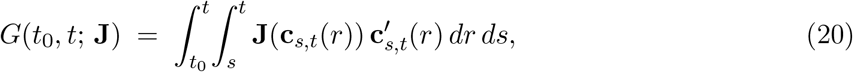

*and* **c**_s,t_ : [*s, t*] → ∝ ^d^ *is any smooth curve with* **c**_s,t_(*s*) = **x**(*s*) *and* **c**_s,t_(*t*) = **x**(*t*).

*Proof*. By the fundamental theorem of calculus along the trajectory,

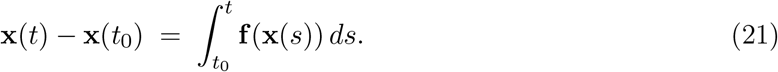

By the row-conservativity path-integral identity, for each *s* ∈ [*t*_0_, *t*],

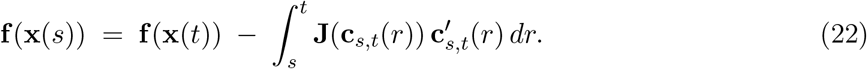

Substituting Eq. 22 into Eq. 21 and pulling **f** (**x**(*t*)) outside the now-trivial *s*-integral yields

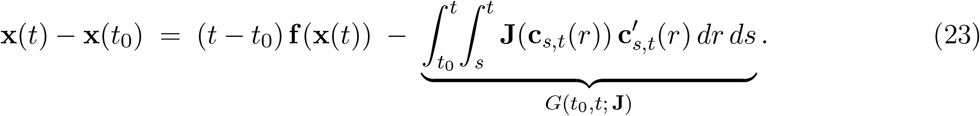

Rearranging gives **f** (**x**(*t*)) = **x**(*t*) − **x**(*t*_0_) + *G*(*t*_0_, *t*; **J**) */*(*t* − *t*_0_).

##### Remark

Equation 19 is implemented in JacobianODE/jacobians/jacobianODE.py: the double integral *G* uses a straight-line path **c**_s,t_(*r*) = **x**(*s*) + (*r* − *s*)(**x**(*t*) − **x**(*s*))*/*(*t* − *s*) with the trapezoid rule from `torchquad` (N=20 subdivisions); the resulting estimate of **f** (**x**(*t*)) is then advanced one step by a standard ODE integrator (default: RK4 from `torchdiffeq`). Because ℱ depends only on **J**, all gradients backpropagate through **Ĵ**^θ^, and no separate **f** -network is trained.

#### B.2 Invariance of the pairwise reachability Gramian trace under isometric direct-sum diffeomorphism

The pairwise reachability Gramian quantifies how strongly source area *β* can drive target area *α* along the reference trajectory. We show that when the encoder respects the subsystem product structure—that is, each subsystem’s latent coordinates depend only on that subsystem’s observed coordinates—and is an isometry on each subsystem factor, the pairwise reachability Gramian computed in the latent is conjugate by an orthogonal map to the Gramian one would compute on the (unknown) data side. Hence the trace and the full eigenvalue spectrum are exactly preserved.

##### Setup

Let 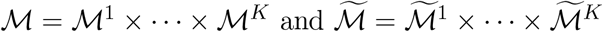 be Riemannian product manifolds with product metrics 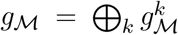 and 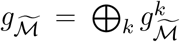 . Assume the encoder is a *direct sum of diffeomorphisms*, with each area’s encoded coordinates depending only on that area’s observed coordinates—this is precisely the per-subsystem encoder construction of the Diffeomorphic-encoder subsection of the main text Methods section,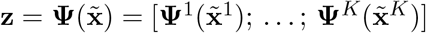,

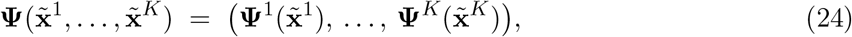

with each **Ψ**^k^ an isometric diffeomorphism. Then **Ψ** is itself an isometry, and its differential is block-diagonal:

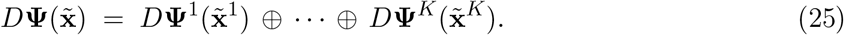

We will write 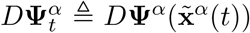 for short, and similarly for the other subsystems. All adjoints (·)^∗^ below are taken with respect to the metric on each side. We write 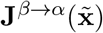 for the dataside cross-area Jacobian block (rows of area *α*, columns of area *β*), 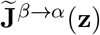 for its latent-side counterpart, **Φ**^α^(*t*_1_, *τ*) for the within-*α* state-transition operator under **J**^α*→*α^, and 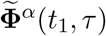 for its latent counterpart. Following the pairwise tangent-space equation in §2,

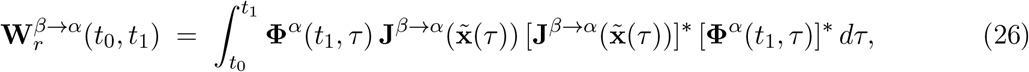

and analogously 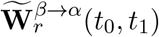 with the tilde objects.

###### Theorem 1

(Spectral invariance of the pairwise reachability Gramian). *Under the setup above, for any pair* (*α, β*) 1, …, *K* ^2^ *with α* ≠ *β, the pairwise reachability Gramian on the latent is conjugate to the pairwise reachability Gramian on the data by an orthogonal map at the final time:*

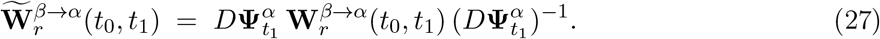

*In particular*,

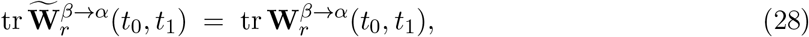

*and the full eigenvalue spectrum—including λ*_min_*—is preserved*.

*Proof*. Each **Ψ**^k^ is an isometry, so *D***Ψ**^k^ is orthogonal:

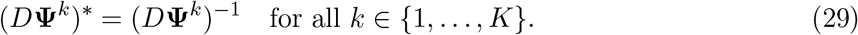

The block-diagonal structure of Eq. 25 means that differentiating 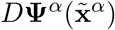 with respect to 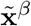 for *β* ≠ *α* gives zero, so the chain rule applied to the cross-area block of the Jacobian yields a clean conjugation *between* the two subsystem factors, with no *D*^2^**Ψ** correction term:

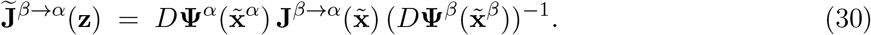

Taking the adjoint of Eq. 30 and using orthogonality,

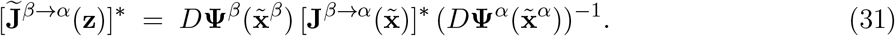

So the *β*-side factors collapse in the symmetric product:

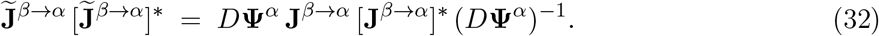

The within-*α* state-transition operator transforms similarly: from 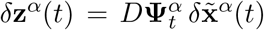 and the definition of **Φ**^α^,

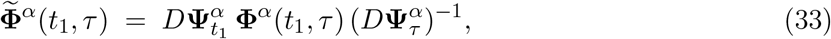

and analogously for the adjoint 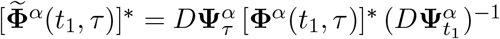. Substituting Eq. 32 and Eq. 33 into the integrand of 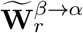,

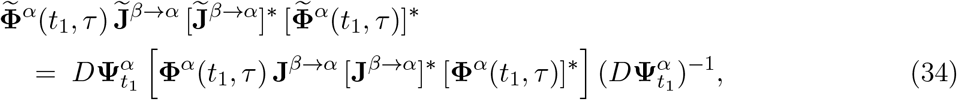

where every intermediate 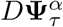 pair cancels: each 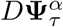 appears next to its inverse, and likewise for the pair that hid inside the symmetric product Eq. 32. Integrating over *τ* and pulling the *τ* -independent outer factors out gives Eq. 27. Trace and eigenvalues are invariant under conjugation by an invertible operator.

*Remark*. The proof extends pointwise to any number of subsystems *K* ≥ 2, with the only requirement being that the encoder respect the subsystem product structure (each **Ψ**^k^ acts only on 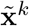). It also extends to the controllability Gramian by replacing **Φ**^α^(*t*_1_, *τ*) with **Φ**^α^(*t*_0_, *τ*) throughout. The hypothesis that each **Ψ**^k^ is an isometry is stronger than what the NICE–Cayley encoder of §C.1 guarantees by construction (the encoder is volume-preserving but not in general isometric); we therefore present the reachability and controllability log-trace summaries reported in the main text as invariants up to encoder-induced similarity, which is what their trace summaries measure. The metric-adjusted Gramian construction of §C.4.3 provides one route to mitigate the residual mismatch by re-weighting the input matrix by the inverse encoder Jacobian on the source-area input space.

#### B.3 Bounded error under generalized teacher forcing

Generalized teacher forcing (GTF)^97^ stabilizes recursive prediction by partially anchoring the rollout to the true trajectory. The strength of anchoring is a scalar *α* ∈ [0, 1]: the input to the next-step network at time *t* is the convex combination

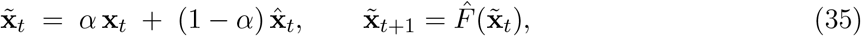

where **x**_t_ is the true state, 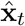 is the (potentially drifting) recursive prediction, and 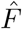is the learned one-step update. *α* = 1 recovers full teacher forcing; *α* = 0 recovers free-running prediction.

##### Lemma 1

(Asymptotic bound on the product of one-step generators). *For every ϵ >* 0 *there exists C* ∈ ℝ_+_ *such that for all T* ≥ 1,

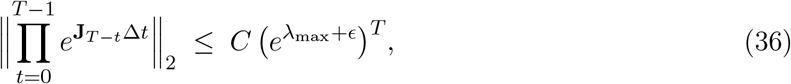

*where* 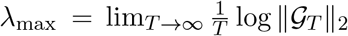 *is the maximum Lyapunov exponent of the linearized system along the trajectory and* 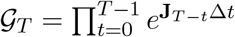.

*Proof*. By the definition of the maximum Lyapunov exponent, for every *ϵ >* 0 there exists *N* such that for all *T > N*, 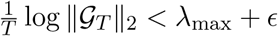, which rearranges to

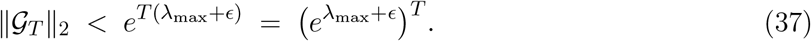

For the pre-asymptotic regime 1 ≤ *T* ≤ *N*, the sequence consists of finitely many bounded matrix products, so there exists *C* ∈ ℝ_+_ such that 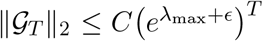 for all 1 ≤ *T* ≤ *N* . Taking the larger of the two constants establishes the claim.

##### Proposition 2

(Bounded error under GTF; cf.^97^). *Let* 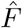 *be the learned one-step update with bounded one-step approximation error*, 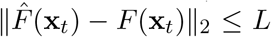 *for some L* ∈ ℝ_+_, *and let the GTF strength satisfy*

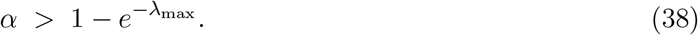

*Then the prediction error* 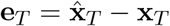 *remains bounded for all T*.

*Proof*. Define 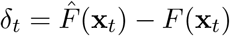 (one-step model error). Note that

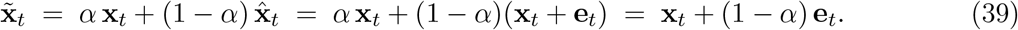

The error propagates as

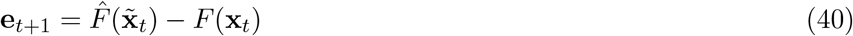

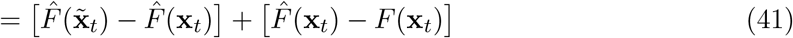

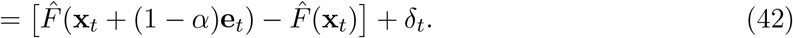

Linearizing 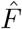 about **x**_t_ (which is valid for sufficiently small **e**_t_),

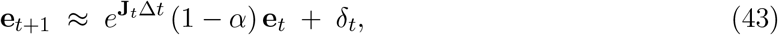

where 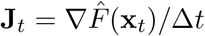 in the continuous-time interpretation (or simply the one-step linearization in the discrete formulation). This recursion has solution

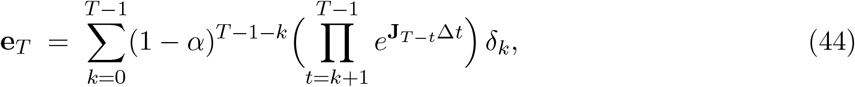

with norm bound

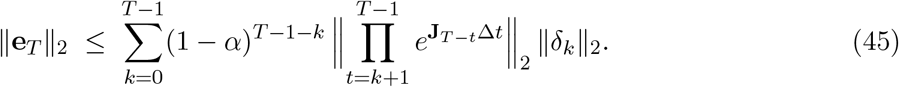

Using ∥*δ*_k_∥_2_ ≤ *L* and Lemma 1,

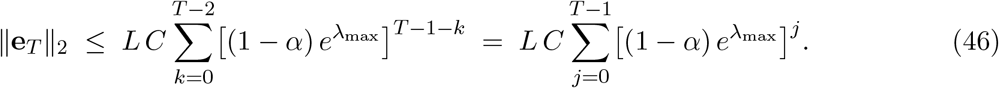

The geometric series on the right converges as *T* → ∞ iff 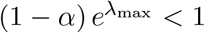, which is equivalent to Eq. 38. Under this condition, 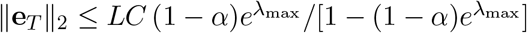 uniformly in *T* .

##### Remark

The bound is tight in the sense that for 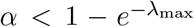 the geometric series diverges and the error grows exponentially; the threshold 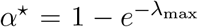 is therefore the minimum GTF strength that ensures bounded recursive prediction in the linearized regime. In practice we use *α*_teacher forcing_ ∈ [*α*_min_, 1] annealed over training (§C.2.2); both the per-rollout maximum effective *λ*_max_ (via the learned Jacobians along the reference trajectory) and the geometric series prefactor enter directly into the validity of the bound.

### C Modeling details

#### C.1 Diffeomorphic encoder

##### C.1.1 Delay embedding

For each subsystem *k* ∈ {1, …, *K*}, the per-subsystem observation 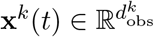 is stacked with *nd* delayed copies at unit stride (*τ* Δ*t* = Δ*t*):

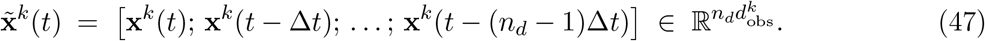

The full delay-embedded observation is the concatenation 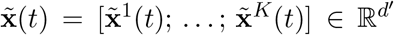 with 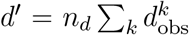, ordered most-recent-first within each per-subsystem block. The choice of *n*_d_ is a per-session sweep variable for the neural data. In practice, we take *τ* Δ*t* = Δ*t* which avoids the dimensionality blow-up of larger strides. For the propofol cohort, *n*_d_ ∈ {5, 10, 15} was searched per session and per area (the per-area block size is *n*_d_ · 64 for the typical 64-channel block).

##### C.1.2 Additive coupling layers

Each additive coupling layer^100^ acts on **y** ∈ ℝ ^n^ by an even/odd channel split **y** = (**a, b**) with **a, b** ∈ ℝ ^n/2^ and applies

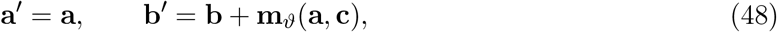

where 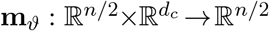 is a conditioner MLP and **c** is a (per-trial, per-batch) condition vector (binary {−1, +1} for awake-vs-anesthesia). The layer Jacobian is

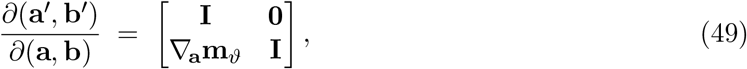

which is unit-triangular: det = 1 identically, regardless of **m**_ϑ_, **a**, or **c**. The layer is exactly invertible by **a** = **a**^*′*^, **b** = **b**^*′*^ − **m**_ϑ_(**a**^*′*^, **c**) in one forward pass.

##### C.1.3 Cayley orthogonal mixers

Between consecutive coupling layers we insert a learnable rotation **Q** ∈ SO(*n*) parameterized by the Cayley transform of a learnable skew-symmetric matrix **A** = −**A**^⊤^ ∈ ℝ^n*×*n^:

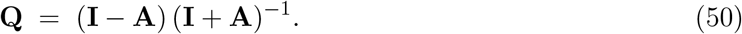

###### Lemma 2

*For any skew-symmetric* **A** *with* −1 ∈*/* spec(**A**), **Q** *is orthogonal with* det **Q** = +1.

*Proof*. **Q**^⊤^**Q** = (**I** + **A**)^*−*⊤^(**I** − **A**)^⊤^(**I** − **A**)(**I** + **A**)^*−*1^ = (**I** − **A**)^*−*1^(**I** + **A**)(**I** − **A**)(**I** + **A**)^*−*1^ = **I** since (**I** − **A**) and (**I** + **A**) share an eigenbasis (commute) and cancel pairwise. The eigenvalues of **A** are purely imaginary in conjugate pairs ± *iω*_k_, so the eigenvalues of **Q** are (1 − *iω*_k_)*/*(1 + *iω*_k_)—complex of unit modulus in conjugate pairs—and their product is +1.

Implementation uses `torch.linalg.solve` (I−A, I+A) rather than an explicit inverse for stability. The skew parameter is built from the strict upper triangle of a free parameter **U** ∈ ℝ ^n*×*n^ via **A** = striu(**U**) − striu(**U**)^⊤^ (the lower triangle and diagonal of **U** are unused, and the operation projects to the skew subspace automatically).

##### C.1.4 Volume-preserving composition and identity-at-initialization

Let 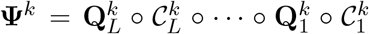 denote the per-subsystem encoder (composition of *L* additive coupling layers 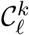 with Cayley mixers 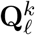). By the chain rule and Lemma 2,

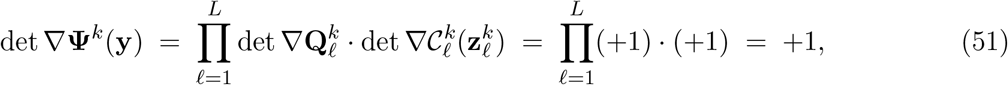

so | det ∇**Ψ**^k^| ≡ 1 everywhere on its domain, and each **Ψ**^k^ is a *C*^∞^ diffeomorphism on 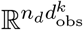. The full encoder **Ψ** = (**Ψ**^1^, …, **Ψ**^K^) inherits these properties on 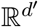 and respects the subsystem product structure required by Theorem 1.

*Identity-at-initialization*. We zero-initialize the last linear layer of every conditioner MLP, which makes **m**_ϑ_(,) ≡**0** at step 0; every coupling layer is then the identity map. We zero-initialize every skew parameter **U** = **0**, which gives **Q** = **I**. A final fixed permutation is appended whose composition with the random per-layer permutations equals the identity, so the entire encoder reduces to **Ψ** = id at step 0. Training therefore departs from the trivial identity-reconstruction solution only as the trajectory and loop-closure losses begin to drive learning, which empirically gives the LR schedule a clean reconstruction-floor starting point.

##### C.1.5 Zero-padded reconstruction loss

The encoder is dimension-preserving on 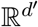, but the intrinsic dimensionality of the dynamics is much smaller. The latent is partitioned as 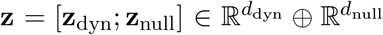 with *d*_null_ = *d*^*′*^ − *d*_dyn_. The dynamic-subspace dimension *d*_dyn_ is fixed per session by a PCA on the delay-embedded training data at variance threshold 0.99 (per-area block selection; the resulting 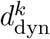 values are concatenated to define the projection 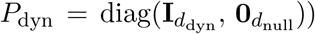. All decoder calls in the model use the zero-padded latent [*P*_dyn_**z**; **0**] rather than **z** itself, so any information that lives in **z**_null_ is ignored on the round-trip. The reconstruction loss (Eq. 10) therefore takes the form

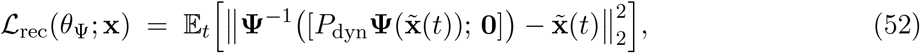

penalizing the round-trip error under the zero-padded forward map. Since **Ψ** is volume-preserving, every information-bearing dimension of the input *must* survive somewhere in **z**; the dynamic/null split together with the zero-padded round-trip squeeze incentivizes the encoder to concentrate the dynamics-relevant information into **z**_dyn_ and dump dynamics-irrelevant content (noise, off-manifold variability) into **z**_null_, where it cannot interfere with the Jacobian-MLP fit.

##### C.1.6 Induced Riemannian metric on the latent

The encoder **Ψ** pulls the (assumed-Euclidean) data-side metric on 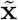 forward to a Riemannian metric on **z**:

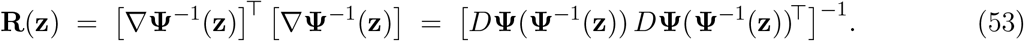

Equivalently, in encoder-Jacobian form, 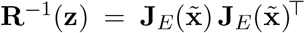 with **J**_E_ = *D***Ψ** evaluated at 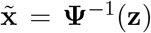. Volume preservation gives det **R** ≡ 1, so the induced metric has unit determinant everywhere; isotropy of **R** (i.e., **R** ≡ **I**) corresponds exactly to the encoder being a Riemannian isometry, which is the hypothesis of Theorem 1. In practice, while we don’t enforce isometry, we find that volume preservation is sufficient to reproduce relative changes in the Gramians on the synthetic systems.

##### C.1.7 Encoder geometry diagnostics: rotation angle, Haar reference, and geodesic distance

Two scalar diagnostics summarize how strongly the trained encoder rotates and warps the latent geometry (Figure 4B,C).

##### Per-layer rotation angle

Each Cayley mixer 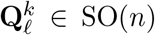 (§C.1.3) has eigenvalues in unit-modulus conjugate pairs 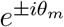 with *θ*_m_ = 2 arctan *ω*_m_, where ±*iω*_m_ are the purely imaginary eigenvalues of its skew generator 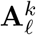. We summarize the rotation of a single mixer by the mean over modes of the absolute eigen-angle,

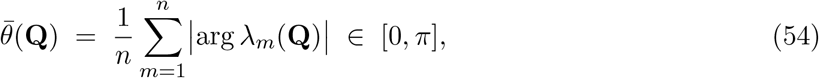

computed numerically as the mean of |angle(eigvals(**Q**)) | ;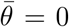 is the identity (no mixing). Figure 4B reports this per Cayley mixer index *ℓ* through encoder depth, both for the individual mixers 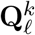 and for the cumulative composite rotation 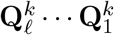 (the net rotation accrued up to depth *ℓ*), averaged over the per-area blocks *k*.

##### Haar SO(*n*) reference

To calibrate the rotation-angle scale we compare against the angle a *maximally mixed* rotation would produce: a matrix drawn uniformly from SO(*n*) under the Haar measure. We sample **G** ∼ N (0, 1)^n*×*n^, take its QR decomposition **G** = **QR**, fix the column signs by **Q** ← **Q** diag(sgn(diag **R**)) (which makes the draw exactly Haar-uniform on O(*n*)), and negate the first column whenever det **Q** *<* 0 to land on the SO(*n*) component. The reference is 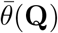 of Eq. 54 averaged over many such draws and over the per-area block dimensions *n*. The trained mixers sit between the identity 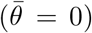 and this Haar saturation level, quantifying how far each layer has rotated relative to a fully randomized rotation of the same dimension.

##### Geodesic distance of the induced metric to the identity

The induced metric **R**(**z**) of §C.1.6 (equivalently its inverse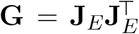) is symmetric positive definite (SPD). The natural distance between SPD matrices is the affine-invariant Riemannian (geodesic) distance, whose value relative to the identity is

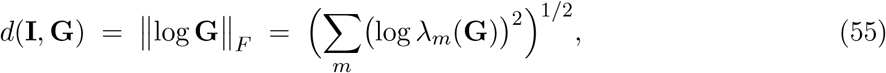

where *λ*_m_(**G**) are the eigenvalues of **G** and log is the matrix logarithm; this is invariant to the choice of **R** versus **R**^*−*1^ = **G** since log **R**^*−*1^ = − log **G**. Writing 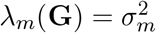 in terms of the singular values *σ*_m_ of the composite encoder Jacobian **J**_E_ along the *d*_dyn_ retained (dynamic-subspace) modes, log *λ*_m_ = 2 log *σ*_m_. We report the per-mode root-mean-square normalization of Eq. 55 (dividing the Frobenius norm by 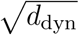 so the scale is *O*(1) in log-stretch units),

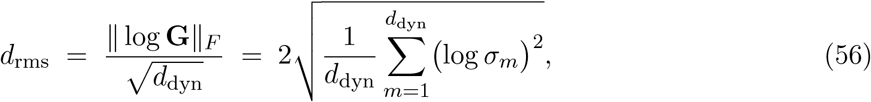

averaged over trajectory samples within each session and condition. *d*_rms_ = 0 exactly when all *σ*_m_ = 1, i.e. when **J**_E_ is orthogonal and the encoder is a local isometry (the hypothesis of Theorem 1); larger values indicate a more anisotropic, more strongly warped latent metric. Because the encoder is volume-preserving (|det **J**_E_ |≡ 1, so the full set of log-singular-values sums to zero), *d*_rms_ reflects the anisotropy of the metric—the spread of its singular values—rather than an overall change of scale. Figure 4C reports the per-session *d*_rms_, awake vs. anesthesia.

#### C.2 JacobianODE-parameterized dynamics

##### C.2.1 Generating predicted states with the Jacobian

The latent JacobianODE rollout is generated as follows. Given an observed conditioning window of length *T*_init_ (Methods: *T*_init_ = 15), the latent states **z**(*t* − (*T*_init_ − 1)Δ*t*), …, **z**(*t*) are encoded once and held fixed. The endpoint of this window, **z**_base_ ≜ **z**_dyn_(*t*), serves as the *base point* of the rollout, and is the only point at which we invoke the double-integral functional of Proposition 1. Concretely, we compute once

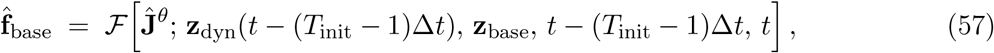

with the double integral *G* (Eq. 20) approximated by the trapezoid rule on the straight-line path between window endpoints (`torchquad`, *N* = 20 subdivisions). All subsequent time-derivative evaluations along the rollout are then obtained by a *single* path integral from this base point, using the row-conservativity identity (Eq. 5):

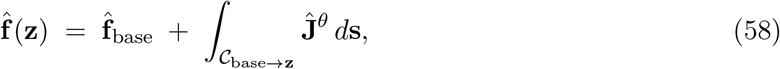

where *C*_base*→***z**_ is the straight-line path from **z**_base_ to **z** (same trapezoid quadrature, *N* = 20). Concretely, at each prediction step *s* = 1, …, *T* :

1. Take the current predicted latent 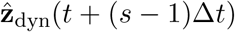 (projected to the dynamic subspace if needed).
2. Evaluate the time derivative 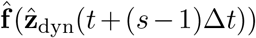 via Eq. 58: a single line integral of **Ĵ**^θ^ from **z**_base_ to the current state, added to the cached 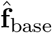 .
3. Advance the latent one step by the RK4 integrator from `torchdiffeq`, producing 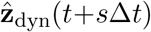.

The number of prediction steps is *T* . We use *T* = 30 for most sample systems except for the partially observed task-trained RNN, for which we use *T* = 20 due to trajectory length constraints. For the neural data, we use *T* = 10 due to the challenging nature of the prediction task. The cumulative latent rollout 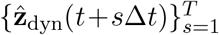 is fed into the latent and decoded prediction losses below. The base point **z**_base_ and its cached derivative 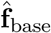 are *not* refreshed during the rollout: the double integral of Proposition 1 is paid exactly once per rollout, and every per-step time-derivative evaluation costs only one straight-line **Ĵ**^θ^ rather than a nested double integral. All gradients still backpropagate through *θ* because 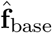 itself is a functional of **Ĵ**^θ^ .

##### C.2.2 Generalized teacher forcing

The recursive rollout is partially anchored to the encoded ground-truth via the GTF input

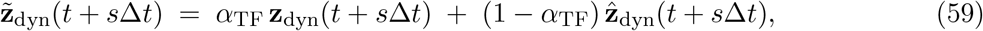

which then feeds the next-step prediction 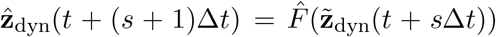. The mixing weight *α*_TF_ is annealed during training from *α*_init_ = 1 (pure teacher forcing) downward toward the *theoretical bound* of Proposition 2,

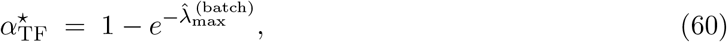

where 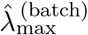 is the maximum Lyapunov exponent of the model’s own linearized rollout, computed per training batch from the predicted Jacobians **Ĵ**^θ^ (**z**(*t*_k_)) along each batch trajectory by the same discrete QR iteration used in §C.5. Because 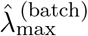 is recomputed at every batch from the current model and the current batch’s reference trajectories, the annealing target is *data-adaptive*: as the model learns more chaotic dynamics, 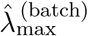 grows, the bound 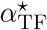 rises toward 1, and the schedule keeps *α*_TF_ above the Proposition 2 threshold that guarantees bounded recursive error. Conversely, when the inferred dynamics are nearly stable, 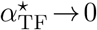 and the schedule permits aggressive annealing toward free-running prediction. The per-batch decay rate (`gamma_teacher_forcing`, default 0.999) controls how quickly *α*_TF_ approaches the current bound from above; a hard floor *α*_min_ = 10^−3^ is enforced for numerical safety so that *α*_TF_ never drops below this value even when the data adaptive bound would permit it. At validation time, the trajectory val_loss is reported at *α*_TF_ = 0 (free-running) and the one-step MASE is reported at *α*_TF_ = 1 (teacher-forced).

##### C.2.3 Latent prediction loss

The latent prediction loss (Eq. 11) penalizes the deviation between the rolled-out latent and the encoded ground-truth latent at each prediction step:

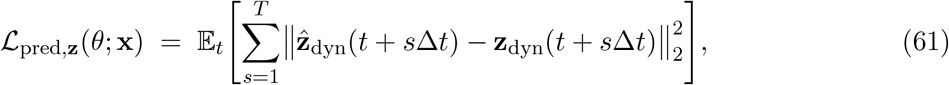

where 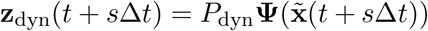 is the encoded ground-truth dynamic latent. In production we use this plain mean-squared error directly, without per-batch generalized-variance normalization or any other rescaling: because the encoder is volume-preserving (§C.1.4), the latent and the delay-embedded observation share an absolute scale, and the reconstruction, latent-prediction, and decoded-prediction MSE terms can be summed at equal weight (§C.6) without further normalization.

##### C.2.4 Decoded prediction loss

Eq. 12 introduces the decoded prediction loss, which compares the decoded prediction against the observed delay-embedded data, projected to the most-recent slice *P*_recent_:

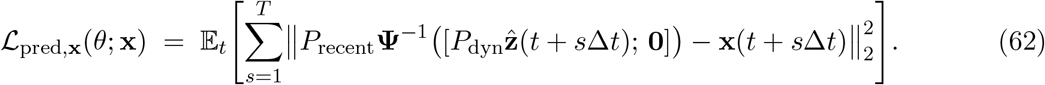

*P*_recent_ extracts the most-recent *d*_obs_ entries of the delay-embedded vector, eliminating redundant comparison against entries that overlap with the conditioning window. This loss closes the loop between the latent dynamics and the data on which they were defined: the encoder is forced to produce a latent whose forward time evolution decodes back to a trajectory that matches the observation. Without this loss the encoder can drift to a latent on which the Jacobian-MLP fits well but whose decoder image is unconstrained.

#### C.3 Loop-closure loss

##### C.3.1 Implementation

Loop-closure regularizes **Ĵ**^θ^ to be row-conservative, exploiting the fact that the path integral of a gradient field along any closed contour vanishes. We sample loops in the dynamic-latent space as concatenations of random straight-line segments connecting randomly selected data points: for each batch we sample *N*_loops_ loops of *N*_loop pts_ ≥ 3 data points each, closing the polygon by appending the first point to the end. The line integral along each segment is approximated by the same trapezoid rule used for the path integral *G* (`torchquad`, *N*_interp_ = 20 subdivisions per segment), and the regularizer is the squared 2-norm of the accumulated row vector across the loop:

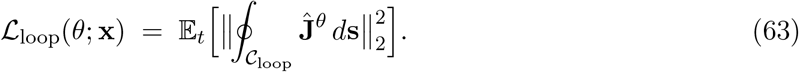

By default the loops are mixed across trajectories within a batch (`mix_trajectories=True`); turning this off restricts each loop to data points from a single trajectory, which we found to be slightly less effective at sampling tangent-space directions orthogonal to the local flow.

##### C.3.2 Consistency under diffeomorphism

A natural concern is that loop closure is defined in the latent rather than the data space. The following proposition verifies that the regularizer is well-posed in the sense that row-conservativity transfers across *C*^1^ diffeomorphisms; in particular, training the latent Jacobian network against this loss is a consistent inductive bias, not a misspecified one.

###### Proposition 3

(Diffeomorphic transfer of row-conservativity). *Let* **f** : ℳ→ ℝ^d^ *be a C*^1^ *vector field on an open set* ℳ⊆ ℝ^d^ *with Jacobian* **J** = ∇**f**, *and let* 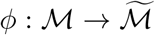 *be a C*^1^ *diffeomorphism with Jacobian* **D** = ∇*ϕ. Define the pushforward field* 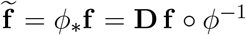 *on* 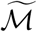 . *Then for every i, the i-th row of* 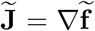 *is the gradient of a scalar potential on* 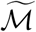 *iff the corresponding row of* **J** *is on* ℳ

*Proof*. On simply connected domains, a *C*^1^ vector field is the gradient of a scalar potential iff its line integral along every closed contour vanishes (Poincaré lemma). Let 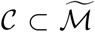 be a closed loop and let 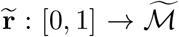 parameterize it. Pull back via *ϕ*^*−*1^ to a loop 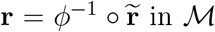 inℳ . For row *i*, with **J**_i,:_ = ∇*u*_i_ for some scalar *u*_i_ : ℳ →ℝ,

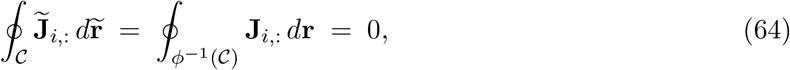

because the line integral of a gradient field along any closed contour vanishes; conversely, every closed loop in ℳ has an image loop in 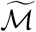 under *ϕ*, so the vanishing transfers in both directions.

#### C.4 Gramian computation

##### C.4.1 Continuous-time definitions

Along a reference trajectory of the linearized subsystem *δ***ż**^α^(*t*) = **J**^α*→*α^(**z**(*t*)) *δ***z**^α^(*t*)+**J**^β*→*α^(**z**(*t*)) *δ***z**^β^(*t*) of the pairwise tangent-space equation in §2, let **Φ**^α^(*t*_2_, *t*_1_) denote the within-target state-transition operator (propagator of *δ***ż**^α^ = **J**^α*→*α^ *δ***z**^α^ on [*t*_1_, *t*_2_]), take the cross-area input matrix as **B**(*τ*) ≜ **J**^β*→*α^(**z**(*τ*)), and take the output matrix as **C**(*τ*) ≜ **J**^α*→*β^(**z**(*τ*))—the *target-to-source* Jacobian block, i.e., the reverse-direction coupling through which the target’s state propagates back into the source’s dynamics. The three pairwise Gramians on [*t*_0_, *t*_1_] are then

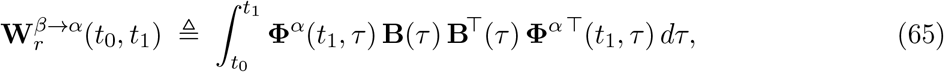

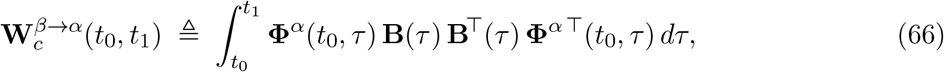

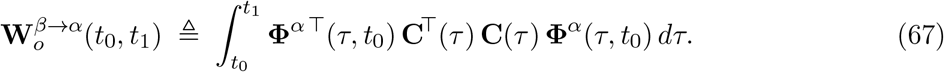

The *reachability* Gramian 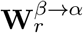 (Eq. 65) measures how much of target subsystem *α*’s state space at the final time *t*_1_ can be reached from a unit-energy input applied through source subsystem *β* over [*t*_0_, *t*_1_]; tr **W**_r_ summarizes the total reachable volume, and *λ*_min_(**W**_r_) the worst-case direction. The *controllability* Gramian 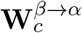 (Eq. 66) is the same input–output coupling propagated to the initial time *t*_0_; equivalently, tr **W**c^*−*1^ is the minimum input energy required to drive *α* from a unit-norm state at *t*_0_ to the origin at *t* . The *observability* Gramian 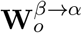 (Eq. 67) measures how strongly perturbations to *α*’s state at *t*_0_ are read off by *β* over [*t*_0_, *t*_1_] via the target-to-source coupling **C**—i.e., how well source area *β* observes target area *α*’s activity. As with reach, the *β* → *α* superscript denotes the actor-on-target direction (here, *β* observing *α*); it is the natural dual of 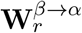 along the same reference trajectory. We do not report observability summaries in the main text but include the definition for completeness because JacobianODE/control/gramians.py accepts (**A, B, C**) = (exp(**J**^α*→*α^Δ*t*), **J**^β*→*α^, **J**^α*→*β^) and computes all three Gramians simultaneously by the same square-root QR mechanics.

##### C.4.2 Square-root QR iteration and numerical stability

We compute the time-varying reachability, controllability, and observability Gramians along the reference trajectory by a low-rank square-root QR iteration^94^, which preserves rank across time steps and keeps the Gramian spectrum reliably computable even when the dynamic range spans many orders of magnitude (as it does for chaotic latents). Implementation lives in `JacobianODE/control/gramians.py` .

Discretize the LTV system along the trajectory at step Δ*t*, **A**_k_ = exp(**J**^α*→*α^(**z**(*t*_k_))Δ*t*), **B**_k_ = **J**^β*→*α^(**z**(*t*_k_)). The square-root factor **S**_r,k_ of the reachability Gramian at step *k* is initialized to zero and updated by

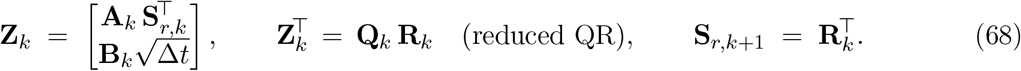

The reachability Gramian at horizon *T* is recovered as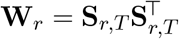 ; the controllability Gramian is computed by the analogous backward sweep with 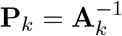 replacing **A**_k_.

##### Why log scaling is required

For chaotic latents the dynamic range of **S**_r,k_ spans many orders of magnitude: the Frobenius norm grows or shrinks at rate 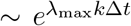 . Over the 10 ms horizon used for the main-text Gramian summaries the per-trajectory factor is modest, but over the longer FTLE/asymptotic horizons used in §C.5 (*T* ∼ 10^4^–10^5^ samples) the unscaled ∥ **S**_r,k_ ∥_F_ leaves the float64 representable range entirely. The eigenvalues of **W**_r_ inherit this dynamic range *squared*, so naively computing log tr **W**_r_ or log *σ*_i_(**W**_r_) on the unrescaled factor produces inf/nan, or quietly saturates at the smallest representable singular value, long before the underlying signal has saturated.

##### The log-prefactor decomposition

We avoid the overflow by representing the square-root factor as a product of an explicit log-domain scalar and a bounded matrix:

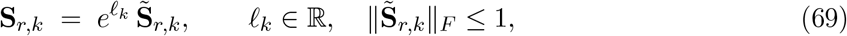

where *ℓ*_k_ accumulates the exponential magnitude and 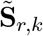 is the rescaled factor whose Frobenius norm is bounded by 1 by construction. The Gramian itself satisfies

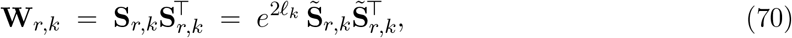

so all eigenvalue and trace summaries factor into a closed-form *ℓ*_k_ contribution plus a bounded contribution from 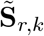.

##### Per-step update

At step *k* we form the stacked matrix

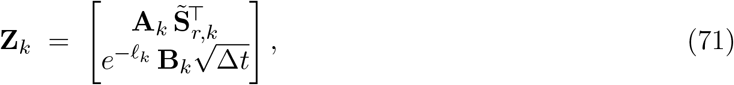

where the driving term is itself rescaled by 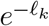 so that both blocks share the same log-domain reference. We then compute *ρ*_k_ = max (∥ **Z**_k_ ∥_F_, *ϵ*) (floored at *ϵ* = `torch.finfo(float64) tiny` to avoid taking the log of zero on initial steps where 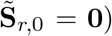, divide **Z**_k_ by *ρ*_k_, reduced-QR-factor the result, and accumulate log *ρ*_k_ into *ℓ*:

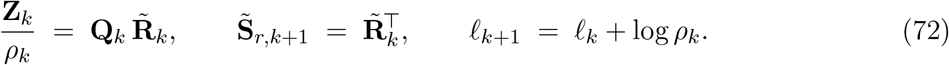

Substituting Eq. 69 into the Gramian recurrence 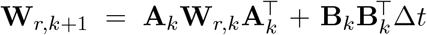 and using 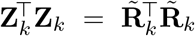 verifies that Eq. 72 preserves the invariant: 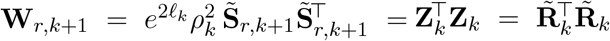 . The bound 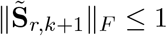 follows from 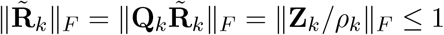.

##### Recovering log-trace and per-mode log-eigenvalues

At horizon *T* we recover the summaries used in the main text (§3) from the pair 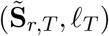. The log-trace is

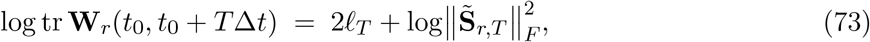

and the per-mode log-eigenvalues come from the SVD 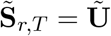 diag 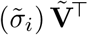:

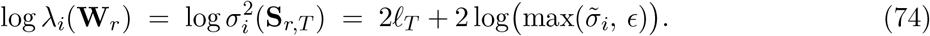

The inverse-log-trace log tr 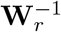 is computed entirely in the log domain via the log-sum-exp identity on the per-mode log-eigenvalues, so the smallest eigenvalues (which dominate tr 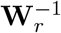 but are most vulnerable to underflow) are recovered without intermediate exponentiation.

##### Numerical safeguards

The full square-root QR iteration runs in `float64`; no intermediates are downcast. The SVD on the rescaled factor uses `torch.linalg.svd` with `driver=`‘gesvd’ (the more numerically stable QR-based driver, preferred over ‘`gesdd`’ for the ill-conditioned rescaled factors that arise near the smallest singular values), and singular values are clamped at *ϵ* before taking logarithms. The log-prefactor floor *ρ*_k_ ≥ *ϵ* guarantees that *ℓ*_k_ stays in the `float64` range even on initial steps where 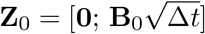 would otherwise have a vanishingly small Frobenius norm.

The horizon used for the main-text propofol analyses is *t*_1_ − *t*_0_ = 10Δ*t* at Δ*t* = 1 ms, i.e., a 10 ms window.

##### C.4.3 Metric-adjusted Gramians

Theorem 1 guarantees that the pairwise reachability Gramian is preserved under an isometric direct-sum encoder. The NICE–Cayley encoder is volume-preserving but in general not isometric, so we additionally provide a metric-adjusted variant of the Gramian (`JacobianODE/control/gramians_metric.py`) that re-weights the input matrix **B** by an arbitrary per-step right-factor **B**_factor_(*τ*):

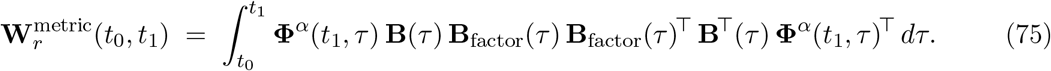

The natural choice is 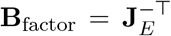 evaluated on the source-area input space at *τ*, which gives 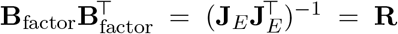, the induced latent-space metric (§C.1.6). Under this choice, control effort is measured in the obs-space-induced inner product on the latent control space rather than in the bare latent Euclidean inner product, which removes the residual non-isometry of the encoder from the reachability calculation. When the factors are absent (**B**_factor_ = **I**), the metric-adjusted routine reproduces the standard square-root QR iteration of §C.4.2 numerically. The metric-adjusted variant is available in the codebase as an experimental construction; in the present paper, the production propofol-cohort figures use the standard square-root QR Gramian of §C.4.2 (no metric adjustment), so the directional reach/ctrl log-trace summaries we report are invariants up to encoder-induced similarity rather than exact diffeomorphism invariants.

#### C.5 Lyapunov exponent computation

The Lyapunov spectrum of the latent dynamics is computed by discrete QR iteration^144^ on the sequence of per-step generators **A**_k_ = exp(**Ĵ**^θ^ (**z**(*t*_k_))Δ*t*). Implementation lives in `JacobianODE/jacobians/analysis`.

Starting from 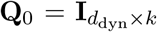 for top-*k* exponents (or *k* = *d*_dyn_ for the full spectrum), at each step we QR-factor

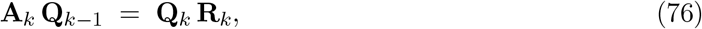

enforcing positive diagonal of **R**_k_ by sign correction (column-wise multiplication of **Q**_k_ by the sign of diag(**R**_k_)), and accumulate the log-diagonal:

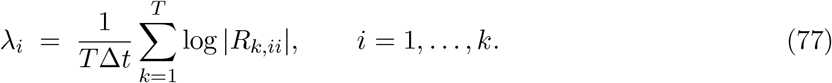

The full spectrum reported in the main text (§3) is estimated per resting-state window (§D.3) and averaged across the windows of each condition: for each window we run the QR iteration over its valid latent trajectory, truncated to *T*_max_ = 2000 samples (2, s at the 1, kHz step, by which the leading exponents have stabilised). Each condition retains ∼ 20–30 windows (∼ !3 × 10^4^ latent samples per condition in aggregate). The exponents are sorted in descending order at the end and divided by Δ*t* to convert to continuous-time exponents. All arithmetic is in float64.

The 10-ms finite-time Lyapunov (FTLE) summaries are derived from the same per-step QR contributions, post-processed in the analysis pipeline by accumulating ∑ _i_ log|*R*_k,ii_|over non-overlapping 10-sample windows and averaging across windows to obtain the per-window generator log-determinant trace. The FTLE-max and FTLE-full-spectrum-mean reported in the main text (§3) are computed at this 10-sample window length.

#### C.6 Training details

##### C.6.1 Optimizer and weight decay

Training uses AdamW^145^ with *β*_1_ = 0.9, *β*_2_ = 0.999, *ε* = 10^−8^, weight decay 10^−4^, and gradient clipping at *ℓ*_2_ norm 1.0. Encoder and Jacobian-MLP parameters share a single optimizer with the same hyperparameters.

##### C.6.2 Learning rate schedule

The learning rate is coupled to the teacher-forcing strength via the `TeacherForcingLRScheduler` (`JacobianODE/jacobians/lightning_base.py`). Let *α* = *α*_TF_ and let 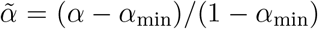 be the renormalized TF strength on [0, 1]. The per-step learning rate is

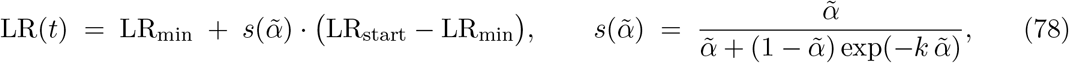

where *k* is a fixed shape parameter. In production we use *k* = 1 (`k_scale: 1` in every `conf/experiment/*.yaml`), which makes 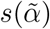 approximately linear in 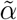 for 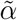 near 1 and slightly compresses LR decay near 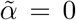 relative to the bare linear schedule *k* = 0; the effect is a more gradual LR drop early in training (when *α*_TF_ is still close to 1) and a slightly steeper drop toward the floor late in training. The two strengths anneal together: as *α*_TF_ decays from *α*_init_ = 1 toward *α*_min_, the LR follows it down toward LR_min_. We use LR_start_ = 10^−4^, LR_min_ = 10^−6^, *α*_min_ = 10^−3^.

##### C.6.3 Early stopping and two-stage protocol

Training runs until early stopping on the free-running trajectory validation loss with patience 5 epochs and a minimum of 15 epochs. For the sample-systems experiments (Lorenz fully observed, Lorenz partially observed, WMTask RNN fully observed, WMTask RNN partially observed), we use the two-stage protocol of^87^ as implemented in `JacobianODE/jacobians/tuning/two_stage_cull.py`: Stage A trains every cell of the sweep grid for 20 epochs (`stage_a_epochs=20`), after which a culling step retains the top- ⌈ (1− *c*)*N*⌉ cells ranked by best-so-far median trajectory val_loss, and Stage B warm-starts the retained cells from the Stage A `last.ckpt` and continues until the early-stopping criterion fires. For the propofol cohort, we ran only a single full sweep—without a Stage A cull—after pilot runs established that the best *n*_d_ always fell within {5, 10, 15 }; the full 3 × 7 per-session grid is then small enough that the cull does not deliver wall-time savings comparable to the sample-systems regime.

#### C.7 Hyperparameter selection

##### C.7.1 Sweep grid

For the propofol cohort, each session is trained on a per-session grid over two hyperparameters: the delay-embedding depth *n*_d_ and the loop-closure regularizer weight *λ*_loop_. The *n*_d_ range{ 5, 10, 15 } was established by pilot runs (early sweeps with a wider candidate set always selected one of these three values); the *λ*_loop_ range is the canonical seven-point geometric grid {0, 10^−6^, …, 10^−1^ } used in^87^. All other architecture and training settings (Table S2) are held fixed across the cohort. The per-session grid therefore has 3 × 7 = 21 cells, and the cohort comprises 21 sessions, for 441 production training runs in total.

**Table S1.**
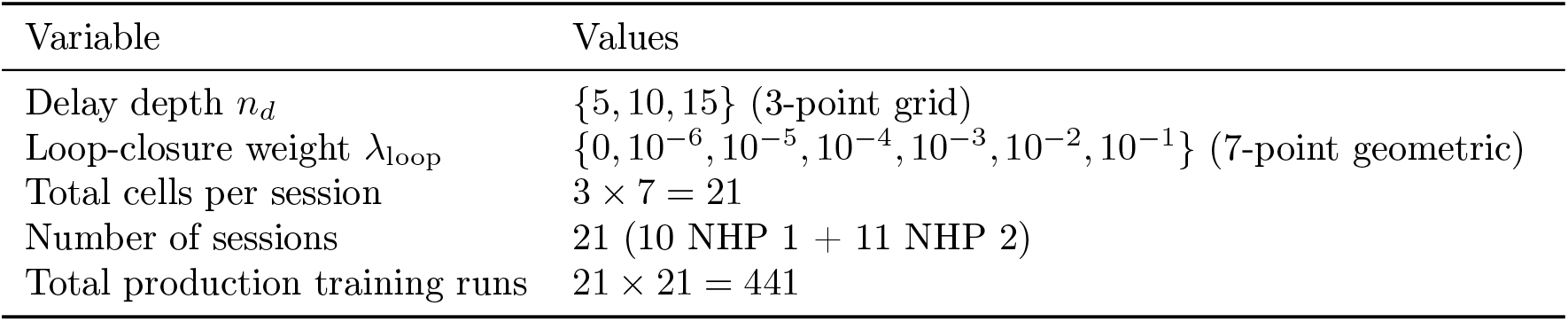
Per-session sweep grid for the propofol cohort. Sample-systems experiments swept a wider grid in a two-stage protocol; see Table S4 for the chosen sample-systems runs.

Fixed per-session architecture and training settings are reported in Table S2.

**Table S2.** Fixed per-session architecture and training settings.

|  |  |
| --- | --- |
| Encoder coupling layers $L$ | 16 |
| Conditioner MLP width | 1024 |
| Conditioner depth | 2 hidden layers, SiLU activation |
| Dynamic dim chosen by | PCA on delay-embedded train, variance threshold 0.99 |
| Jacobian MLP widths | (2048, 2048, 4096, 4096), SiLU activation |
| Condition vector dim $d_c$ | 1 (binary $\{-1, +1\}$) |
| Latent prediction loss weight | 1 |
| Decoded prediction loss weight | 1 |
| Reconstruction loss weight | 1 |
| Trajectory sequence length | 25 samples |
| Sequence spacing | 15 samples |
| Prediction steps | 10 |
| Initial window $T_{\text{init}}$ | 15 |
| Training noise (`obs_noise_scale`) | 0 for the chosen production cohort |
| Optimizer | AdamW, weight decay $10^{-4}$ |
| LR schedule | TeacherForcingLRScheduler, $10^{-4} \rightarrow 10^{-6}$ |
| Early stopping patience | 5 epochs (free-running trajectory val_loss) |
| Minimum epochs | 15 |

##### C.7.2 Selection criteria

For each session we select a single chosen run from the 21-cell sweep by applying two hard criteria (C1 and C2, Table S3) and ranking the survivors by free-running trajectory validation loss. The criteria gate the cells before ranking rather than serving as a post-hoc tiebreaker, so a cell that achieves the lowest trajectory val_loss but fails C1 or C2 is rejected. C1 (decoder-corrected one-step MASE *<* 1) certifies that the model’s one-step prediction beats the decoded-persistence baseline on the validation split; C2 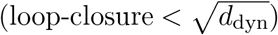 certifies that the learned Jacobian is approximately row-conservative on the per-session dynamic subspace. We do not gate on C3 (small fast-eigenvalue fraction): although it is computed alongside C1 and C2 in `JacobianODE/jacobians/tuning/selection.py`, dropping it from the selection rule produces the identical chosen run on every session of the propofol cohort, so it does not bind in practice. The implementation reads the per-cell val metrics from the W&B logger, evaluates each criterion, and returns the chosen run name (which we recover from `chosen_extras.pkl` for downstream analysis).

**Table S3.** Per-session model-selection criteria. Among the 21 cells of each session’s sweep, the chosen run is the one that minimizes the free-running trajectory val_loss subject to passing both criteria.

|  |  |
| --- | --- |
| C1 (one-step accuracy) | decoder-corrected one-step MASE $< 1$ on the validation split. The decoder-corrected variant subtracts the reconstruction floor by comparing the predicted next-step against the *decoded encoded ground-truth next-step* rather than the raw observation; this prevents the reconstruction floor itself from disqualifying otherwise-good cells. |
| C2 (loop closure) | loop-closure validation loss $< \sqrt{d_{\text{dyn}}}$, with $d_{\text{dyn}}$ the per-session dynamic-subspace dimension. |
| Tie-break | among cells passing C1–C2, the one minimizing free-running trajectory val_loss is chosen. |

##### C.7.3 Sample-systems run identifiers and chosen hyperparameters

The W&B run identifiers for the four sample-systems runs reported in the main text (§3) are reported in Table S4.

**Table S4.**
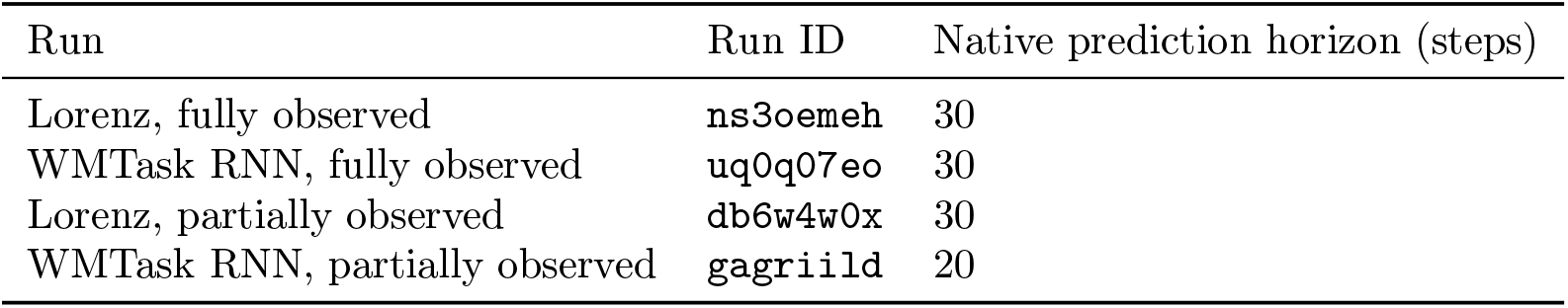
Run IDs for the four sample-systems runs reported in the main text (§3).

| Run | Run ID | Native prediction horizon (steps) |
| --- | --- | --- |
| Lorenz, fully observed | ns3oemeh | 30 |
| WMTask RNN, fully observed | uq0q07eo | 30 |
| Lorenz, partially observed | db6w4w0x | 30 |
| WMTask RNN, partially observed | gagriild | 20 |

Evaluation is on the held-out test split at 10 prediction steps with *α*_TF_ = 0 (free-running) for *R*^2^ and *α*_TF_ = 1 (teacher-forced) for one-step MASE. Selection criteria match the per-session criteria above (C1–C2 with best-trajectory-loss tie-break).

##### C.7.4 Per-session chosen hyperparameters for the propofol cohort

The chosen *n*_d_ landed at each of {5, 10, 15 }at least twice across the cohort, and *λ*_loop_ ranged from 10^−5^ to 10^−1^; no single sweep cell dominated, confirming that the per-session selection reflects genuine cross-session variation rather than a degenerate winner. Any analysis directories carrying the `smallenc` recipe slug (a separate encoder-size investigation that is not part of the production cohort) are excluded from this table at the extraction stage; the audit CSV emitted by the extraction script documents the exclusions.

**Table S5.** Per-session chosen hyperparameters under the C1–C2 criterion across the 21 propofol-cohort sessions. The chosen (*n*_*d*_, *λ*_loop_) pair was selected from each session’s 3 × 7 sweep by minimizing the free-running trajectory val_loss subject to C1–C2; *d*_dyn_ is the per-session PCA-99% dynamic-subspace dimension on the delay-embedded train data (sum across the four areas), and depends on *n*_*d*_. Session labels follow the `<YYYYMMDD>-<NN>` convention. Values extracted from the on-disk `chosen_extras.pkl` and per-session `report.md` artifacts; see `figures/paper/extract_chosen_hyperparams.py`.

| Session | Animal | $n_d$ | $\lambda_{\text{loop}}$ | $d_{\text{dyn}}$ |
| --- | --- | --- | --- | --- |
| 20160809-01 | NHP 1 | 5 | $10^{-4}$ | 115 |
| 20160818-02 | NHP 1 | 5 | $10^{-2}$ | 93 |
| 20160822-02 | NHP 1 | 5 | $10^{-1}$ | 97 |
| 20160826-02 | NHP 1 | 10 | $10^{-3}$ | 129 |
| 20160831-02 | NHP 1 | 15 | $10^{-2}$ | 96 |
| 20160902-02 | NHP 1 | 10 | $10^{-4}$ | 135 |
| 20160908-02 | NHP 1 | 5 | $10^{-2}$ | 128 |
| 20160912-02 | NHP 1 | 10 | $10^{-3}$ | 144 |
| 20160914-02 | NHP 1 | 10 | $10^{-1}$ | 136 |
| 20160916-02 | NHP 1 | 5 | $10^{-3}$ | 110 |
| 20160105-01 | NHP 2 | 10 | $10^{-1}$ | 176 |
| 20160107-01 | NHP 2 | 10 | $10^{-3}$ | 146 |
| 20160109-01 | NHP 2 | 10 | $10^{-2}$ | 134 |
| 20160113-01 | NHP 2 | 10 | $10^{-3}$ | 153 |
| 20160121-01 | NHP 2 | 5 | $10^{-4}$ | 107 |
| 20160123-01 | NHP 2 | 10 | $10^{-5}$ | 133 |
| 20160125-01 | NHP 2 | 10 | $10^{-4}$ | 164 |
| 20160201-01 | NHP 2 | 10 | $10^{-4}$ | 173 |
| 20160206-01 | NHP 2 | 5 | $10^{-1}$ | 120 |
| 20160210-01 | NHP 2 | 15 | $10^{-5}$ | 166 |
| 20160301-01 | NHP 2 | 5 | $10^{-1}$ | 128 |

### D Experimental details

#### D.1 Dynamical systems data

##### D.1.1 Lorenz system

We use the canonical Lorenz^103^ system, 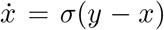, 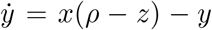, *ż* = *xy* − *βz*, with the standard chaotic parameters *σ* = 10, *ρ* = 28, *β* = 8*/*3. Following^87^, trajectories are simulated using the `dysts` package^146^, which samples each dynamical system at 100 time steps per characteristic Fourier-spectrum timescale. Integration uses `scipy.integrate.solve_ivp` (default `Radau`, or `RK45` when configured) through the `dysts` backend (`JacobianODE/dysts_sim/flows.py`). We use *n*_ics_ = 32 trajectories of *n*_periods_ = 12 Fourier periods each (1200 samples per trajectory), with initial conditions perturbed from the system’s default IC by Gaussian noise of standard deviation 0.2 (`new_ic_mode=`‘`random`’, `traj_offset_sd=0.2`). The train/test split is the standard 80/20 split across trajectories. For the fully observed variant, we expose all three dimensions (*x, y, z*) as the per-subsystem observation. For the partially observed variant, we observe only the *x* coordinate (*d*_obs_ = 1) and delay-embed before encoding. To evaluate the methods under observation noise, we additionally train with 1%, 5%, and 10% Gaussian observation noise added i.i.d. over time, where the percentage is the ratio of the Euclidean norm of the noise to the mean Euclidean norm of the data.

##### D.1.2 Task-trained working-memory selection RNN

The 128-dimensional task-trained RNN is the working-memory selection network of^87^, inspired by^104^ (codebase: `ControlJacobians/wmtask/wmtask.py`). The RNN has continuous-time dynamics

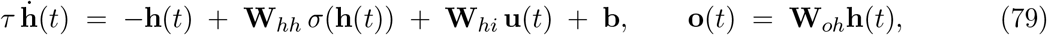

discretized in implementation by Euler integration as **h**_k+1_ = **h**_k_+(Δ*t/τ*) −**h**_k_+(**W**_hi_⊙mask_in_)**u**_k_+ Whh σ(**h***k*) + b with τ = 0.05 s and Δt = 0.02 s. Here **W**_hh_ ∈ ℝ^128*×*128^ defines the recurrent weights, **W**_hi_ maps the input into the hidden state, **W**_oh_ ∈ ℝ^4*×*128^ maps the hidden state to a four-dimensional output, **b** is a bias term, and *σ* is the exponential linear unit (ELU) activation (*σ*^*′*^(*h*) = 1 for *h >* 0 and *σ*^*′*^(*h*) = exp(*h*) otherwise). The network has two 64-neuron areas (denoted *N*_1_ = *N*_2_ = 64 in the codebase): a “visual” area (which receives sensory input via an input mask zeroing the cognitive-area inputs) and a “cognitive” area (whose readout mask zeros the visual-area outputs). The within-area weights are initialized with greater connectivity strength than the across-area weights; because input enters only the visual area and output exits only the cognitive area, the two areas are forced to interact to solve the task. The RNN solves a working-memory selection task with trial structure (in milliseconds): fixation 500, stimuli 500, delay 1 750, cue 100, delay 2 750, response 250 (total 2850 ms); on each trial two of four colors are presented (one-hot vectors) during the stimulus epoch, and the cue then selects one of the two for the response. The RNN achieves 100% accuracy. JacobianODE training is performed on the post-cue delay + response portion of the trial, where the dynamics are state-locked to the cued color. The discrete Jacobian along the rollout is **J**_disc_ = **I** + (Δ*t/τ*) (− **I** + **W**_hh_ diag(*σ*^*′*^(**h**))). For the partially observed variant, we retain 16 randomly chosen channels from each of the two areas (*d*_obs_ = 32 total). We condition the model on the discrete trial-condition variable **c** ∈ {−1, +1} corresponding to the cued color. As in the Lorenz case, we add 5% Gaussian observation noise during training;^87^ reports that this improves Jacobian estimation across all systems considered there.

##### D.1.3 Synthetic two-area linearization (conceptual control factors)

Figure 1E illustrates, on a small synthetic linear system, how the pairwise reachability and controllability Gramians depend on four properties of a source → target interaction: the target area’s dynamic stability, and the magnitude, alignment, and dimensionality of the cross-area coupling block (codebase: `JacobianODE/_notebook/render_damping_vs_coupling.py`). In every case we form a source → target control system (**A**_tgt_, **B, C**)—with **A**_tgt_ the target’s within-area dynamics and **B** = **C** the coupling block playing the role of **J**^β*→*α^—and compute the reachability and controllability Gramians by the same square-root QR pipeline used for the learned models (§C.4.2), over a horizon of *T* = 1000 steps at Δ*t* = 0.01 (10 s), reporting log tr **W**.

##### Stability and magnitude (coupled damped-oscillator areas)

For the stability and magnitude factors the system is an 8-D pair of two 4-D area s. Each] area’s 4 × 4 intrinsic block is a direct sum of two planar damped-rotation blocks 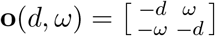 at frequencies *ω*_1_ = 1.0 and *ω*_2_ = 2.3 and common damping *d*; the coupling is isotropic, . To vary *dynamic stability* we sweep the target damping *d* = *b* from 0.8 down to 0.02 (40 log-spaced values; source damping fixed at 0.8, *g* = 0.25); the abscissa is the system stability − max_i_ Re *λ*_i_(**A**), which decreases as *b* → 0. To vary *coupling magnitude* we sweep *g* from 0.02 to 1.5 (50 linearly-spaced values; target damping fixed at 0.4), so that ∥**B**∥_F_ = 2*g*.

##### Alignment and dimensionality (fixed diagonal target, shaped coupling)

For the alignment and dimensionality factors we fix a 4-D target with diagonal dynamics **A**_tgt_ = −diag(0.3, 0.6, 1.0, 1.5) (eigen-timescales ordered slow→fast) and shape the coupling block **B** to have a prescribed set of squared singular values 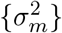 with left singular vectors aligned to the target eigenbasis, holding the total energy fixed at 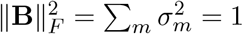 *Coupling alignment* is varied by sweeping the energy 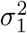 that **B** places on the slowest target mode from 0.05 to 0.95 (30 values), spreading the remaining energy uniformly over the other modes. *Coupling dimensionality* is varied by sweeping the participation ratio 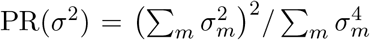 of the coupling spectrum (from ≈ 2 to ≈ 3) along two mirrored paths that redistribute the off-leading energy toward either the slowest or the fastest minor mode; the two paths share the same PR at each step but yield different Gramians, illustrating that the dimensionality scalar alone does not determine reachability/controllability—the direction in which the coupling spreads its energy, relative to the target’s own timescales, also matters.

#### D.2 Non-human primate electrophysiology

The neural data for this study is the propofol cohort previously deposited in association with^92^, comprising 21 sessions from two rhesus macaques (*Macaca mulatta*), denoted NHP 1 and NHP 2. NHP 1 was female, aged 8 years, ∼ 6.6 kg; NHP 2 was male, aged 14 years, ∼ 13.0 kg. All procedures followed the guidelines of the Massachusetts Institute of Technology Committee on Animal Care and the National Institutes of Health. The animals were pair-housed in temperature-controlled environments, were not involved in previous procedures, and were surgically implanted with a subcutaneous vascular access port (Model CP-6, Norfolk Access Technologies, Skokie, IL) at the cervicothoracic junction of the neck, with the catheter tip reaching the termination of the superior vena cava via the external jugular vein. Both NHPs were implanted with 8 × 8 chronic Utah arrays (“Utah arrays”, MultiPort: 1.0 mm shank length, 400 *µ*m spacing, Blackrock Microsystems, Salt Lake City, UT), yielding 64 channels per multi-electrode array. Electrodes were placed in four areas: ventrolateral prefrontal cortex (vlPFC), frontal eye fields (FEF), posterior parietal cortex (PPC), and auditory cortex (STG). In each session, the NHP performed a non-demanding pre-anesthesia task (passive airpuff/tone classical conditioning); following this task, the NHP was infused with propofol anesthesia for 60 minutes (30 minutes at a higher loading dose of 0.58 mg kg^*−*1^ min^*−*1^ for NHP 1 and0.285 mg kg^*−*1^ min^*−*1^ for NHP 2; 30 minutes at a lower maintenance dose of 0.32 mg kg^*−*1^ min^*−*1^ for NHP 1 and ∼ 0.075 mg kg^*−*1^ min^*−*1^ for NHP 2). The infusion was then stopped and the NHP gradually returned to a normal awake state. Propofol was intravenously infused via a computer-controlled syringe pump (PHD ULTRA 4400, Harvard Apparatus).

##### LFP recording and preprocessing

Local field potentials were recorded at 30 kHz and filtered online via a lowpass at 250 Hz, then downsampled to 1 kHz. Areas STG and PPC shared a common subdural ground/reference; areas FEF and vlPFC each had their own ground/reference. Across recording sessions, a synchronization test signal with locally unique temporal structure was simultaneously injected on one auxiliary analog channel of each system and used to rectify residual cross-system timing offsets (event-code shifts plus linear interpolation to a common time base). Following^92^ we removed the across-session per-channel mean from each LFP, line-noise-removed via temporally windowed sinusoidal fits at 60 Hz and all harmonics (and at the empirically found 107.35, 214.7, 190.2, 196.8, 393.6 Hz line-noise frequencies), and low-pass filtered each LFP at 300 Hz with a third-order bidirectional Butterworth. Signals were subsequently low-pass filtered at 80 Hz. We did not remove eye-movement or muscle artifacts: intracortical recordings (in contrast to scalp EEG/MEG) are not subject to the same scalp-contact contamination, so these signals are non-cerebral by location but not by amplitude (the recording site is below the dura). All four areas were included in the analysis with no per-channel averaging within an area: the analysis models population dynamics, and channel averaging discards the information that the per-channel representation transforms over time.

##### Amplitude-threshold artifact rejection

To remove residual recording artifacts (e.g. electrical transients or movement-induced glitches) we applied the same amplitude-threshold rejection procedure used in the cross-anesthetic destabilization study of Eisen et al. [93] (the same procedure was introduced for the propofol cohort in^92^). We first split each session into non-overlapping windows of the length used for analysis (*W* = 15 s; §D.3). Within each window we computed, for every electrode, its maximum absolute amplitude over the full window. If two or more electrodes exceeded 1 mV within a window, that window was excluded from all subsequent analysis; any electrode that exceeded the 1 mV threshold was additionally excluded from the remainder of that session. As an added safeguard against onset transients, we also excluded the window immediately preceding any window in which an artifact was detected. Across the propofol cohort this procedure removed between 4 and 23 electrodes per session and between 6% and 44% of windows per session. The surviving per-channel, per-window validity mask is the noise-valid mask consumed by the resting-state window selection of §D.3; because electrodes were never averaged within an area (above), a rejected electrode is simply dropped from the population state for the remainder of the session rather than imputed.

#### D.3 Selection of resting-state windows

Resting-state windows for the propofol cohort are carved by the loader, which retains only the two protocol epochs of interest (“awake” and “maintenance dose”) and explicitly excludes the induction and loading-dose transients (these epochs are simply not assigned to any of the per-condition output sections). Within each retained epoch, candidate windows begin at least *t*_post trial_ = 3.0 s after each preceding behavioral-trial offset, are *W* = 15 s long, and are advanced with stride *S* = 15 s (non-overlapping). A candidate is kept only if at least 1.5 s of contiguous samples survive the per-channel noise-valid mask (computed from a per-session amplitude-threshold rejector); shorter contiguous blocks are coalesced where possible. Each retained window is assigned to a condition by the position of its midpoint relative to infusion start. The default per-channel decimation factor is 1 (no downsampling). Because windows are non-overlapping with stride equal to length, the resulting awake and maintenance corpora are exhaustive on the retained epochs and identical across sessions in structure; the only per-session degrees of freedom are the noise-valid channel mask and the exact number of windows that survive coalescing in each condition.

### E Statistical analyses

#### E.1 Sign-agreement gate

For every 4 × 4 source–target cell or per-area summary axis, we compute the per-NHP mean of the per-session paired difference (anesthesia − awake), separately for NHP 1 and NHP 2. A cell is admitted to significance testing only if these two per-NHP means agree in sign. The gate is a guard against pooled artifacts: a near-zero pooled difference produced by two equal-and-opposite per-NHP shifts is not a population-level signal, and the gate rejects such cells before any *p*-value is computed. We do not interpret cells that fail this gate even if the pooled Wilcoxon would have been significant.

#### E.2 Pooled signed-rank test

For the per-pair 4 × 4 grids we apply a one-sided Wilcoxon signed-rank test in the agreed-upon direction. For the per-area summary axes we apply a two-sided test on the pooled paired differences. For the session-level single-quantity metrics (*R*^2^, decoder-corrected MASE) we report per-NHP and pooled paired two-sided Wilcoxon. Wilcoxon was chosen over a paired *t*-test because the per-session paired differences are heavy-tailed in several metrics and the sample size (*n* = 21) is small enough that the assumptions of the *t*-test cannot be confidently verified. All Wilcoxon tests use `scipy.stats.wilcoxon` with the `zero_method=`‘`wilcox`’ convention for tied pairs.

#### E.3 Benjamini–Hochberg correction across the 4×4 grid

For each 4 × 4 grid (reach trace, reach inverse trace, ctrl trace, ctrl inverse trace, block Frobenius norm, block participation ratio) we apply Benjamini–Hochberg FDR control across the 16 cells at *q <* 0.05. The grid-level family is the appropriate test family because the cells of each grid share a common per-session denominator and the substantive question is which cells of the grid carry the signal. The per-area summary axes are tested separately at *p <* 0.05 because the axis-level family is single-axis.

### F Per-pair statistical tables

The main text reports the 4 × 4 source–target heatmaps of mean differences (anesthesia − awake) for the six per-pair quantities tested in this study and flags significant cells in the figures. The following tables provide the underlying numerical detail for each cell that passes the sign-agree + Benjamini– Hochberg gate at *q <* 0.05 (§E.1–E.3). For each significant cell we list: the source → target pair, the pooled paired-difference mean ± standard error of the mean (across all 21 sessions), the per-NHP mean differences separately for NHP 1 and NHP 2 (which by the sign-agree gate must share the same sign), and the (one-sided, in the agreed direction) pooled Wilcoxon signed-rank *p*-value before BH correction. Cells that fail the sign-agree gate or the BH-FDR cut are omitted from each table. The six tables in this section cover the four directional Gramian summaries reported in Figure 5 (reach log tr **W**, reach log tr **W**^*−*1^, ctrl log tr **W**, ctrl log tr **W**^*−*1^) plus the per-block Jacobian summaries (block Frobenius norm, block participation ratio).

#### Reachability log-trace log tr W

Per-pair (anesthesia − awake) differences in reach log tr **W** are reported in Table S6.

**Table S6.** Per-pair reach log tr **W**, anesthesia awake, *n* = 21 sessions. All cells listed pass the sign-agree + BH-FDR gate at *q <* 0.05.

| source → target | pooled $\Delta \pm \text{SEM}$ | per-NHP $\Delta$ (NHP 1, NHP 2) | $p$ |
| --- | --- | --- | --- |
| PPC → PPC | $-0.1006 \pm 0.0130$ | $-0.1218, -0.0813$ | $9.54 \times 10^{-7}$ |
| PPC → STG | $-0.0709 \pm 0.0328$ | $-0.1228, -0.0237$ | 0.01753 |
| PPC → vIPFC | $+0.2451 \pm 0.0774$ | $+0.3348, +0.1635$ | 0.006346 |
| STG → PPC | $-0.3917 \pm 0.0555$ | $-0.5249, -0.2706$ | $4.77 \times 10^{-7}$ |
| STG → STG | $-0.0666 \pm 0.0127$ | $-0.0824, -0.0522$ | $5.25 \times 10^{-5}$ |
| STG → FEF | $-0.1267 \pm 0.0334$ | $-0.1117, -0.1404$ | 0.0008001 |
| STG → vIPFC | $-0.1179 \pm 0.0275$ | $-0.1089, -0.1261$ | $4.20 \times 10^{-5}$ |
| FEF → PPC | $-0.3001 \pm 0.0325$ | $-0.3361, -0.2673$ | $4.77 \times 10^{-7}$ |
| FEF → STG | $-0.2026 \pm 0.0421$ | $-0.3147, -0.1008$ | $8.06 \times 10^{-5}$ |
| FEF → FEF | $-0.1518 \pm 0.0109$ | $-0.1689, -0.1362$ | $4.77 \times 10^{-7}$ |
| FEF → vIPFC | $+0.1330 \pm 0.0479$ | $+0.1110, +0.1531$ | 0.009737 |
| vIPFC → PPC | $-0.2767 \pm 0.0465$ | $-0.3887, -0.1748$ | $1.43 \times 10^{-6}$ |
| vIPFC → STG | $-0.3110 \pm 0.0375$ | $-0.3667, -0.2604$ | $9.54 \times 10^{-7}$ |
| vIPFC → vIPFC | $-0.1292 \pm 0.0155$ | $-0.1682, -0.0936$ | $1.43 \times 10^{-6}$ |

#### Reachability inverse log-trace log tr **W**^*−*1^

Per-pair (anesthesia − awake) differences in reach log tr **W**^*−*1^ are reported in Table S7.

#### Controllability log-trace log tr W

Per-pair (anesthesia − awake) differences in ctrl log tr **W** are reported in Table S8.

**Table S7.** Per-pair reach log tr **W**^*−*1^, anesthesia − awake. All cells listed pass the gate.

| source → target | pooled $\Delta \pm \text{SEM}$ | per-NHP $\Delta$ (NHP 1, NHP 2) | $p$ |
| --- | --- | --- | --- |
| PPC → PPC | $+0.4176 \pm 0.1347$ | $+0.7230, +0.1400$ | $0.0001464$ |
| PPC → STG | $+0.4167 \pm 0.0669$ | $+0.3490, +0.4782$ | $9.06 \times 10^{-6}$ |
| PPC → FEF | $+0.3425 \pm 0.0681$ | $+0.4447, +0.2497$ | $9.06 \times 10^{-6}$ |
| STG → PPC | $+0.3682 \pm 0.0339$ | $+0.3707, +0.3659$ | $4.77 \times 10^{-7}$ |
| STG → STG | $+0.1855 \pm 0.0557$ | $+0.1444, +0.2228$ | $1.57 \times 10^{-5}$ |
| STG → FEF | $+0.3242 \pm 0.0402$ | $+0.4042, +0.2515$ | $4.77 \times 10^{-7}$ |
| STG → vlPFC | $+0.2791 \pm 0.0278$ | $+0.2322, +0.3217$ | $4.77 \times 10^{-7}$ |
| FEF → PPC | $+0.3394 \pm 0.0335$ | $+0.3027, +0.3727$ | $4.77 \times 10^{-7}$ |
| FEF → STG | $+0.3546 \pm 0.0339$ | $+0.3917, +0.3208$ | $4.77 \times 10^{-7}$ |
| FEF → FEF | $+0.3347 \pm 0.1043$ | $+0.2788, +0.3856$ | $4.77 \times 10^{-7}$ |
| FEF → vlPFC | $+0.2154 \pm 0.0272$ | $+0.2051, +0.2248$ | $4.77 \times 10^{-7}$ |
| vlPFC → PPC | $+0.4035 \pm 0.0397$ | $+0.4622, +0.3501$ | $4.77 \times 10^{-7}$ |
| vlPFC → STG | $+0.4827 \pm 0.0617$ | $+0.4959, +0.4706$ | $2.38 \times 10^{-6}$ |
| vlPFC → FEF | $+0.4442 \pm 0.0747$ | $+0.5966, +0.3056$ | $1.43 \times 10^{-6}$ |

**Table S8.** Per-pair ctrl log trW, anesthesia ™ awake. All cells listed pass the gate.

| source → target | pooled $\Delta \pm \text{SEM}$ | per-NHP $\Delta$ (NHP 1, NHP 2) | $p$ |
| --- | --- | --- | --- |
| PPC → PPC | $-0.3193 \pm 0.0475$ | $-0.3105, -0.3274$ | $3.34 \times 10^{-6}$ |
| PPC → STG | $-0.4217 \pm 0.0527$ | $-0.4447, -0.4008$ | $2.38 \times 10^{-6}$ |
| PPC → FEF | $-0.2975 \pm 0.0578$ | $-0.3062, -0.2895$ | $4.77 \times 10^{-6}$ |
| STG → PPC | $-0.5151 \pm 0.0791$ | $-0.6274, -0.4131$ | $4.77 \times 10^{-7}$ |
| STG → STG | $-0.3830 \pm 0.0427$ | $-0.3511, -0.4121$ | $4.77 \times 10^{-7}$ |
| STG → FEF | $-0.4218 \pm 0.0531$ | $-0.4735, -0.3748$ | $4.77 \times 10^{-7}$ |
| STG → vlPFC | $-0.3410 \pm 0.0343$ | $-0.2940, -0.3838$ | $4.77 \times 10^{-7}$ |
| FEF → PPC | $-0.4867 \pm 0.0706$ | $-0.5920, -0.3909$ | $3.34 \times 10^{-6}$ |
| FEF → STG | $-0.6202 \pm 0.0623$ | $-0.7128, -0.5361$ | $4.77 \times 10^{-7}$ |
| FEF → FEF | $-0.2879 \pm 0.0339$ | $-0.3195, -0.2592$ | $4.77 \times 10^{-7}$ |
| FEF → vlPFC | $-0.1653 \pm 0.0578$ | $-0.1496, -0.1796$ | $0.005063$ |
| vlPFC → PPC | $-0.4950 \pm 0.0655$ | $-0.5774, -0.4202$ | $1.43 \times 10^{-6}$ |
| vlPFC → STG | $-0.6657 \pm 0.0536$ | $-0.6806, -0.6523$ | $4.77 \times 10^{-7}$ |
| vlPFC → FEF | $-0.4487 \pm 0.0637$ | $-0.5770, -0.3320$ | $4.77 \times 10^{-7}$ |
| vlPFC → vlPFC | $-0.2069 \pm 0.0224$ | $-0.2004, -0.2127$ | $4.77 \times 10^{-7}$ |

#### Controllability inverse log-trace log tr **W**^*−*1^

Per-pair (anesthesia − awake) differences in ctrl log tr **W**^*−*1^ are reported in Table S9.

#### Jacobian block Frobenius norm ∥**Ĵ**_ij_ ∥_F_

Per-pair (anesthesia − awake) differences in the Jacobian block Frobenius norm are reported in Table S10.

**Table S9.** Per-pair ctrl log tr **W**^*−*1^, anesthesia − awake. All cells listed pass the gate.

| source → target | pooled $\Delta \pm \text{SEM}$ | per-NHP $\Delta$ (NHP 1, NHP 2) | $p$ |
| --- | --- | --- | --- |
| PPC → PPC | $+0.5058 \pm 0.1475$ | $+0.8585, +0.1853$ | $6.53 \times 10^{-5}$ |
| PPC → STG | $+0.4384 \pm 0.0706$ | $+0.3351, +0.5323$ | $1.19 \times 10^{-5}$ |
| PPC → FEF | $+0.4495 \pm 0.0751$ | $+0.5844, +0.3268$ | $9.54 \times 10^{-7}$ |
| PPC → vlPFC | $+0.1757 \pm 0.0510$ | $+0.1295, +0.2176$ | 0.001876 |
| STG → PPC | $+0.4083 \pm 0.0362$ | $+0.4392, +0.3802$ | $4.77 \times 10^{-7}$ |
| STG → STG | $+0.2374 \pm 0.0583$ | $+0.1913, +0.2794$ | $2.62 \times 10^{-5}$ |
| STG → FEF | $+0.4461 \pm 0.0473$ | $+0.5567, +0.3456$ | $4.77 \times 10^{-7}$ |
| STG → vlPFC | $+0.4118 \pm 0.0283$ | $+0.3845, +0.4366$ | $4.77 \times 10^{-7}$ |
| FEF → PPC | $+0.3786 \pm 0.0363$ | $+0.3637, +0.3922$ | $4.77 \times 10^{-7}$ |
| FEF → STG | $+0.3888 \pm 0.0380$ | $+0.4051, +0.3740$ | $4.77 \times 10^{-7}$ |
| FEF → FEF | $+0.4758 \pm 0.1091$ | $+0.4705, +0.4805$ | $4.77 \times 10^{-7}$ |
| FEF → vlPFC | $+0.3386 \pm 0.0312$ | $+0.3371, +0.3399$ | $4.77 \times 10^{-7}$ |
| vlPFC → PPC | $+0.4511 \pm 0.0496$ | $+0.5531, +0.3583$ | $4.77 \times 10^{-7}$ |
| vlPFC → STG | $+0.5252 \pm 0.0600$ | $+0.5085, +0.5403$ | $9.54 \times 10^{-7}$ |
| vlPFC → FEF | $+0.5602 \pm 0.0801$ | $+0.7317, +0.4044$ | $4.77 \times 10^{-7}$ |

**Table S10.** Per-pair ∥Ĵ ij∥F, anesthesia ™ awake. All cells listed pass the gate.

| source → target | pooled $\Delta \pm \text{SEM}$ | per-NHP $\Delta$ (NHP 1, NHP 2) | $p$ |
| --- | --- | --- | --- |
| PPC → PPC | $-47.4514 \pm 7.7469$ | $-65.5622, -30.9870$ | $1.19 \times 10^{-5}$ |
| PPC → STG | $-25.0390 \pm 2.9885$ | $-30.0683, -20.4668$ | $4.77 \times 10^{-7}$ |
| PPC → FEF | $-18.3192 \pm 1.7879$ | $-19.1833, -17.5336$ | $4.77 \times 10^{-7}$ |
| PPC → vlPFC | $-13.6493 \pm 1.3243$ | $-11.9255, -15.2164$ | $4.77 \times 10^{-7}$ |
| STG → PPC | $-8.1960 \pm 1.6578$ | $-11.8104, -4.9101$ | $5.25 \times 10^{-5}$ |
| STG → STG | $-72.6155 \pm 7.6688$ | $-70.2020, -74.8096$ | $9.54 \times 10^{-7}$ |
| STG → FEF | $-17.3419 \pm 1.9623$ | $-20.6818, -14.3057$ | $4.77 \times 10^{-7}$ |
| STG → vlPFC | $-18.1553 \pm 1.8810$ | $-14.9224, -21.0943$ | $4.77 \times 10^{-7}$ |
| FEF → PPC | $-2.3021 \pm 1.0957$ | $-1.4782, -3.0511$ | 0.03507 |
| FEF → STG | $-13.4517 \pm 1.3642$ | $-13.5055, -13.4027$ | $4.77 \times 10^{-7}$ |
| FEF → FEF | $-118.1231 \pm 9.7532$ | $-138.5780, -99.5278$ | $4.77 \times 10^{-7}$ |
| FEF → vlPFC | $-10.3758 \pm 1.5540$ | $-13.4146, -7.6131$ | $3.34 \times 10^{-6}$ |
| vlPFC → STG | $-14.6888 \pm 1.5594$ | $-9.9267, -19.0181$ | $4.77 \times 10^{-7}$ |
| vlPFC → FEF | $-4.2621 \pm 2.3600$ | $-5.5720, -3.0713$ | 0.04790 |
| vlPFC → vlPFC | $-89.6821 \pm 8.0704$ | $-93.6591, -86.0666$ | $4.77 \times 10^{-7}$ |

#### Jacobian block participation ratio

PR(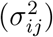)

Per-pair (anesthesia − awake) differences in the Jacobian block participation ratio are reported in Table S11.

### G Per-area Lyapunov: leading exponent and within-area spectra

Per-area within-area Lyapunov leading-exponent results (anesthesia − awake) are reported in Table S12.

**Table S11.** Per-pair PR(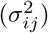), anesthesia − awake. Only six cells pass the gate (four diagonal blocks plus the FEF↔vlPFC reciprocal pair).

| source → target | pooled $\Delta \pm \text{SEM}$ | per-NHP $\Delta$ (NHP 1, NHP 2) | $p$ |
| --- | --- | --- | --- |
| PPC → PPC | $-0.5508 \pm 0.0991$ | $-0.5169, -0.5817$ | $6.68 \times 10^{-6}$ |
| STG → STG | $-0.3202 \pm 0.0926$ | $-0.0498, -0.5661$ | 0.0001206 |
| FEF → FEF | $-0.8093 \pm 0.1111$ | $-0.8368, -0.7842$ | $4.77 \times 10^{-7}$ |
| FEF → vIPFC | $-0.4312 \pm 0.1403$ | $-0.4433, -0.4201$ | 0.0008001 |
| vIPFC → FEF | $-0.6870 \pm 0.1330$ | $-0.7032, -0.6721$ | $1.57 \times 10^{-5}$ |
| vIPFC → vIPFC | $-0.8698 \pm 0.1159$ | $-0.9023, -0.8402$ | $4.77 \times 10^{-7}$ |

**Table S12.** Per-area within-area Lyapunov leading exponent max *λ* (1/s), anesthesia awake. Per-area paired two-sided Wilcoxon; “sign-agree” indicates whether the per-NHP mean differences agree.

| Area | $\max \lambda$ awake | $\max \lambda$ anesthesia | $\Delta$ | Wilcoxon $p$ | sign-agree |
| --- | --- | --- | --- | --- | --- |
| PPC | $-14.33 \pm 2.49$ | $-13.53 \pm 2.97$ | $+0.81 \pm 1.10$ | 0.68 | no |
| STG | $-27.15 \pm 3.73$ | $-25.95 \pm 3.32$ | $+1.20 \pm 0.73$ | 0.14 | yes |
| FEF | $-36.37 \pm 3.44$ | $-34.02 \pm 3.47$ | $+2.35 \pm 0.73$ | 0.0025 | yes |
| vIPFC | $-28.88 \pm 3.48$ | $-30.29 \pm 3.63$ | $-1.40 \pm 1.30$ | 0.29 | no |

The single per-area cell that passes the sign-agree gate on the leading exponent is FEF; the per-area full-spectrum mean (reported in the main text (§3)) passes the gate on STG, FEF, and vlPFC, indicating that the cohort-wide destabilization signal is broad-spectrum rather than concentrated in the leading mode.

### H Per-NHP session bookkeeping

Per-animal session counts and per-condition window lengths are reported in Table S13.

**Table S13.** Session counts per animal and per condition. Resting-state windows: 15 min awake-baseline, 30 min maintenance-dose centered on the middle of the maintenance epoch.

| Animal | sessions | awake (min) | maintenance (min) | areas |
| --- | --- | --- | --- | --- |
| NHP 1 | 10 | 15 | 30 | PPC, STG, FEF, vIPFC |
| NHP 2 | 11 | 15 | 30 | PPC, STG, FEF, vIPFC |
| Total | 21 | — | — | — |

### I Software and reproducibility

All numeric results can be reproduced by checking out the JacobianODE library at commit `4c6e86f8` and the MindControl analysis pipeline at the corresponding commit on the `main` branch, and running the sweep with the hyperparameter grid in Table S1 on the sessions listed in the per-NHP book-keeping table. PyTorch 2.7.1+cu118, torchdiffeq 0.2.5, SciPy 1.17.1 are pinned in the repository’s environment file. Training was carried out on the Engaging compute cluster at MIT.

